# Structural insights into the opening mechanism of C1C2 channelrhodopsin

**DOI:** 10.1101/2024.11.14.623569

**Authors:** Matthias Mulder, Songhwan Hwang, Matthias Broser, Steffen Brünle, Petr Skopintsev, Caspar Schattenberg, Christina Schnick, Sina Hartmann, Jonathan Church, Igor Schapiro, Florian Dworkowski, Tobias Weinert, Peter Hegemann, Han Sun, Jörg Standfuss

**Author notes:** authors contributed equally (Matthias Mulder, Songhwan Hwang).

## Abstract

Channelrhodopsins, light-gated cation channels, enable precise control of neural cell depolarization or hyperpolarization with light in the field of optogenetics. This study integrates time-resolved serial crystallography and atomistic molecular dynamics (MD) simulations to resolve the structural changes during C1C2 channelrhodopsin activation. Our observations reveal that within the crystal environment, C1C2 predominantly remains in a light-activated state with characteristics of the M_390_ intermediate. Here, rearrangement of retinal within its binding pocket partially opens the central gate towards the extracellular vestibule. These structural changes initiate channel opening but were insufficient to allow K^+^ flow. Adjusting protonation states to represent the subsequent N_520_ intermediate in our MD simulations induced further conformational changes, including rearrangements of transmembrane helices 2 and 7, that opened the putative ion-translocation pathway. This allows spontaneous but low cation but not anion conduction, that matches experiments. Our findings provide critical structural insights into key intermediates of the channel opening mechanism, enhancing our understanding of ion conduction and selectivity in channelrhodopsins at an atomistic level.

## Introduction

Channelrhodopsins are light-gated ion channels that allow mobile algae cells to find suitable conditions for photosynthesis. Beside their natural role as sensory photoreceptors, scientists have employed wild-type channelrhodopsins and their variants to control the depolarization of neural cells with light giving birth to the field of optogenetics. Targeting a genetically defined set of neurons can be used to understand their contribution to the behavior of animals^1^ or for medical applications like the partial restoration of vision in Retinitis Pigmentosa human patients^2^. Optogenetic applications have sparked a wide interest into the study of channelrhodopsins to facilitate the design of improved variants with shifted absorption wavelengths^3–5^, reduced desensitization^6^, or modified ion conductance^7–11^. Uncovering the molecular principles of how channelrhodopsins open in response to light is of high biophysical interest and may promote the design of additional optogenetic tools for neurobiology.

The photocycle of the most widely used optogenetic tool, channelrhodopsin-2 (ChR2) from *Chlamydomonas reinhardtii*, is well-known from extensive spectroscopic studies and could be related to time-resolved photocurrent measurements in cellular membranes^1,12–14^. The activation cycle of dark-adapted ChR2 is initiated by the light-induced trans-to-cis isomerization of the trans,15-anti retinyliden chromophore into the first ground state K_520_ intermediate within 2 picoseconds^15^. This principle intermediate further relaxes into an L-type intermediate and a subsequent early M-species with deprotonated chromophore M_390a_. This intermediate, formed within 1 μs, provides the principal signal that initiates partial opening of the channel. However, primary channel opening occurs on a 100 μs timescale which is delayed compared to M_390a_ formation. This led to the suggestion that another conformational change compared to the M1 to M2 transition in bacteriorhodopsin (BR)^16^ is associated with the primary pore opening in a M_390a_- to-M_390b_ transition within about 100 μs^14,17^. This M_390b_-state primarily conducts protons as shown by laser-stimulated electrophysiology^14^. The fully open state with Na^+^ and K^+^ conductance is approached after 1 ms and contains a reprotonated chromophore named N_520_. Channel closing occurs on a 30 ms time scale under standard conditions and in detergent-solubilized protein, which however in cellular conditions strongly varies with extracellular and intracellular pH and membrane voltage. The LA_480_ state was originally assigned as a late photocycle intermediate is in reality a “light-adapted dark state”. It is a result of a C13=C14 and C15-N double isomerization into a 13-cis,15-syn species that only within minutes reconverts to the fully dark-adapted state^14,18–20^. Photoactivation of the light-adapted dark state again initiates C13=C14 isomerization and a syn- cycle with all photointermediates in syn-configuration and a low conductance state with prolongated decay.

The first crystal structure of a channelrhodopsin has been solved for a chimera between Channelrhodopsin-1 and Channelrhodopsin-2 from *Chlamydomonas reinhardtii* commonly known as C1C2^4^. This chimera with transmembrane helices (TM) 1-5 from ChR1 and 6-7 from ChR2 is more stable in detergent than the parental ChRs and provided the first invaluable insight into the overall architecture of the central ion conducting core^4^. It shares the prototypical seven-transmembrane helical fold of microbial ion pumps and human G protein-coupled receptors. The structure further revealed a partial ion-conducting pathway reaching from the retinal binding pocket towards the extracellular side, lined by negatively charged residues suggested to secure cation selectivity. Very similar arrangements have been found in related structures of chrimson^21^ and channelrhodopsin-2^22^. Indeed, some of these residues were later included in the structure-guided design of a light-gated chloride channel^7,10^.

Despite these successes it remains largely unknown how the channel opens at the atomistic scale due to lack of an open state structure. A first constriction of the putative channel, called the central gate which is composed of residues Glu129, Asn297 and Ser102, is found close to the retinal Schiff base and its counterion residues Glu162 and Asp292 (Channelrhodopsin-2 numbering can be obtained by subtracting 39 from the C1C2 residue numbers). A second ion constriction site, called the intracellular gate because of its location closer to the cytosolic site, is formed by Glu122, His173 and Tyr109. Both gates would need to open to connect the intracellular side with the vestibule on the extracellular side of the putative channel to allow full conductance. But the molecular details and sequence of events leading to the open state remain largely elusive.

Time-resolved serial crystallography is a powerful technique to study structural changes in light-activated proteins with up to femtosecond temporal and near-atomic spatial resolution^23^. It has been employed on a number of rhodopsins^24^, including C1C2 where it revealed early rearrangements in the retinal binding pocket^25^. Here we report how we used serial synchrotron crystallographic methods, initially developed to resolve late photointermediates in BR^26^, to study light-activation of C1C2 in the millisecond range. We present conformational movements after photoactivation in the retinal binding pocket, as well as in the adjacent gates and relate these results to channel activation mechanisms. Starting from this structural intermediate, we adjusted the protonation state of the titratable residues involved in channel opening and performed large-scale molecular dynamics (MD) simulations under different transmembrane potentials. These simulations further opened up the C1C2 channel, resulting in a fully open N_520_-like state that revealed a number of cation permeation events, with the simulated K^+^ conductance matching the experimental ranges. Our study provides structural insights into key intermediates of the channel opening during the photocycle, enabling the understanding of ion conduction and selectivity in channelrhodopsins at an atomistic scale.

## Results

### Light-activated structure of C1C2

For our room temperature analysis of light-induced conformational changes in C1C2 we purified recombinant C1C2 from insect cells and grew crystals in lipidic mesophases following established conditions^4^ and subjected them to serial synchrotron crystallography^27^ at the Swiss Light Source, where they diffracted with anisotropic resolution up to 2.6 Å (**Supplementary Table 1)**. An initial dataset collected without illumination resulted in a dark state structure that is principally identical to earlier reports (root means square deviations of Cα atoms = 0.42 and 0.63 to 3UG9^4^ and 7C86^25^, respectively). Next, we collected data while continuously illuminating the extruding lipidic cubic phase with the C1C2 crystals using a laser diode. The speed of extrusion was adjusted so that crystals were illuminated for approximately 100 ms before diffraction patterns were recorded. In comparison to other rhodopsins, retinal isomerization in C1C2 has a relatively low quantum efficiency around 30 percent^28^ and, in addition, single flash illumination of dark-adapted channelrhodopsins can yield a fraction of the protein in the desensitized light-adapted state^14^. Together these effects likely resulted in the low activation levels in a previous time-resolved X-ray Free Electron Laser study relying on nanosecond laser excitation^25^. The millisecond exposure with our laser diode, even at the employed low laser fluence of 0.2 mJ/cm^2^, led to higher activation levels of 30 percent and excellent Fo(light)-Fo(dark) difference electron density maps (**Supplementary Figure Figure 1A)**. Data was of sufficient quality to calculate extrapolated structure factor amplitudes and to model structural changes upon illumination (**Figure 1**).

**Figure 1:**
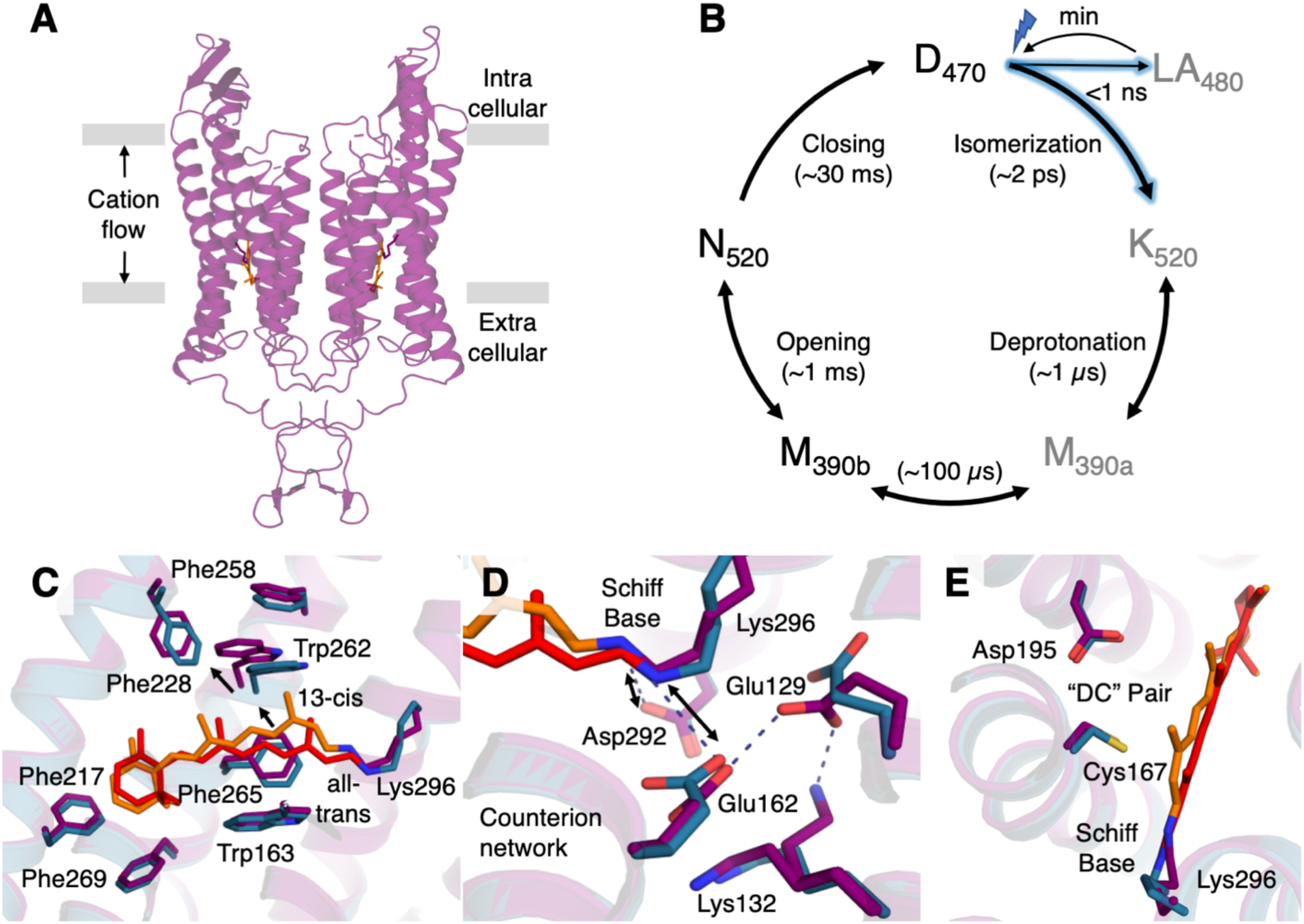
Light-activation of C1C2 channelrhodopsin. (**A**) Overview of the C1C2 dimer structure. The cellular membrane and the main flow of cations are indicated. (**B**) The channelrhodopsin photocycle is composed of several spectroscopic intermediates with characteristic absorption maxima that depend on the environment of the retinal chromophore. The first transition (D_470_-to-K_520_) is dominated by retinal isomerization, followed by Schiff base deprotonation (K_520_-to-M_390_) and reprotonation (M_390_-to-N_520_) during channel opening. Approximate times are shown based on spectroscopic studies in solution. (**C**) The light-activated structure shows how the binding pocket has adapted to retinal isomerization. (**D**) Local rearrangements in the counterion network indicate deprotonation of the retinal Schiff base without (**E**) rearrangements in the DC pair.

In order to further narrow the time where structural changes occur and to reduce the probability of dark-adapted to light-adapted conversion, we have illuminated crystals for 5 ms and collected X-ray data for 200 ms in bins of 5 ms. Pearson correlation analysis (based on^29^ with modifications described in^30^) of electron density changes over time (**Supplementary Figure 1B**) confirm accumulation of the light-activated intermediate within about 50 ms and its subsequent decay towards the dark state structure.

### Effect of retinal isomerization on the overall C1C2 structure

The initiating event in the photocycle of microbial rhodopsin is the all-trans to 13-cis isomerization of the retinylidene molecule. Beyond sharing this fundamental reaction, the conformations do vary between rhodopsins and can furthermore change throughout the photocycle, in BR, for example, from a twisted conformation in the K state towards a fully planarized retinal in the M intermediate^31^. The retinal molecule in our structure of photoactivated C1C2 has fully isomerized into a planar 13-cis conformation that tilts the retinal polyene backbone towards the intracellular side (**Figure 1C**), as well as shifting the chromophore sidewards towards TM3 and TM4. Interestingly, this conformation is markedly different to those observed in the previous X-ray laser study of C1C2 activation^21^ and much closer to that of the BR M-intermediate^32^ (**Supplementary Figure 2**). The binding pocket in C1C2 is lined by aromatic residues (Trp163, Phe217, Trp262, Phe265 and Phe269) that hold the hydrophobic polyene backbone and the β-ionone ring of the retinal in place. Our light-activated structure shows major rearrangements in this hydrophobic pocket (**Figure 1C**). In particular the shift in Trp262, likely resulting from a steric clash with the C20 methyl group of the retinal, is very pronounced and transmits further changes towards the intracellular side of TM5 and TM6. Relocation of the β-ionone ring away from Phe269 and Phe217, together with the shift of the polyene towards Phe265 and away from Trp163 introduce further changes into the hydrophobic core of the protein. Remarkably, these substantial early rearrangements occur with only minor changes in the overall packing of the 7TM bundle (root mean square deviation of Cα = 0.4 Å), similar to what we observed in early intermediates of other rhodopsins^30,31,33^.

### The light-activated structure represents an early deprotonated intermediate

Beside retinal isomerization as the primary trigger of activation in all rhodopsins, the state of the retinylidene that links the retinal chromophore to TM7 is of particular functional relevance. Since channelrhodopsin photoactivation can lead to the formation of a 13-cis,15-anti retinylidene of the standard photocycle and with lower efficiency 13-cis,15-syn retinylidene of the light-adapted state both configurations have to be considered. While both configurations are structurally very similar, refinement using 13-cis-anti restrains yielded the overall better fit to the electron density map.

The second critical step in channelrhodopsin activation is controlled by the protonation state of the Schiff base, which deprotonates in the K_520_-to-M_390_ transition and is reprotonated in the M_390_-to- N_520_ transition^34,35^. In C1C2, two negatively charged residues, Glu162 and Asp292, are close enough to act as counterion to stabilize protonation of the retinal Schiff base^36^. Proton transfer from the Schiff base to Asp292 leads to a strong blue shift of absorption, the hallmark of the M_390_ intermediates which is required for the cation channel to open^12^. In our light-activated C1C2 structure distances between the Schiff base and Glu162 and Asp292 have increased to 5.2 and 3.9 Å, respectively (**Figure 1D)**. While we can’t observe protonation states directly, the breaking of these critical interactions suggests the Schiff base to be deprotonated in our light-activated structure.

A third step is enabled by reprotonation of the Schiff base during the M_390_-to-N_520_ transition with the proton coming most likely from a water around the Schiff base^37^. Mutations in residues Asp195 and Cys167 strongly alter gating kinetics^38^ suggesting high relevance for the transition to the conductive state^39,40^, even though the on-gating function of this pair has never been substantiated. In our light-activated structure the isomerized retinal is shifted towards this “DC pair” and a negative difference peak on Wat4 indicates that it has shifted from its position between Asp195 and Cys167. However, we do not observe the extreme shift of the retinal towards the “DC pair” suggested by previous TR-SFX^25^ experiments and computer simulations^41^. In our light-activated structure, the “DC pair” remains basically in the same position (**Figure 1E)** as in the dark state, confirming that reprotonation of the Schiff base has not occurred yet.

In concert with the changes in the retinal and Glu162, Lys132 adopts an alternative conformation and forms an ionic interaction with the central gate residue Glu129 (**Figure 1D**). Probing the structure with a sphere of 1.6 Å diameter placed into the extracellular vestibule results into a continuous channel through this constricting region towards the intracellular gate formed by Glu122, His173 and Tyr109 (**Figure 2**). These residues adopt principally the same positions as in our structure of the C1C2 dark state indicating, that this gate remains in a closed conformation. Based on these observations, we suggest our light-activated structure to represent a M_390_-like intermediate where the Schiff base is deprotonated and the channel is only partially opened and conductive only for protons. This assignment is in agreement with time-resolved spectroscopy on C1C2 crystals^25^, where N_520_ formation is hindered and the M_390_ intermediate accumulates within milliseconds after photoactivation.

**Figure 2:**
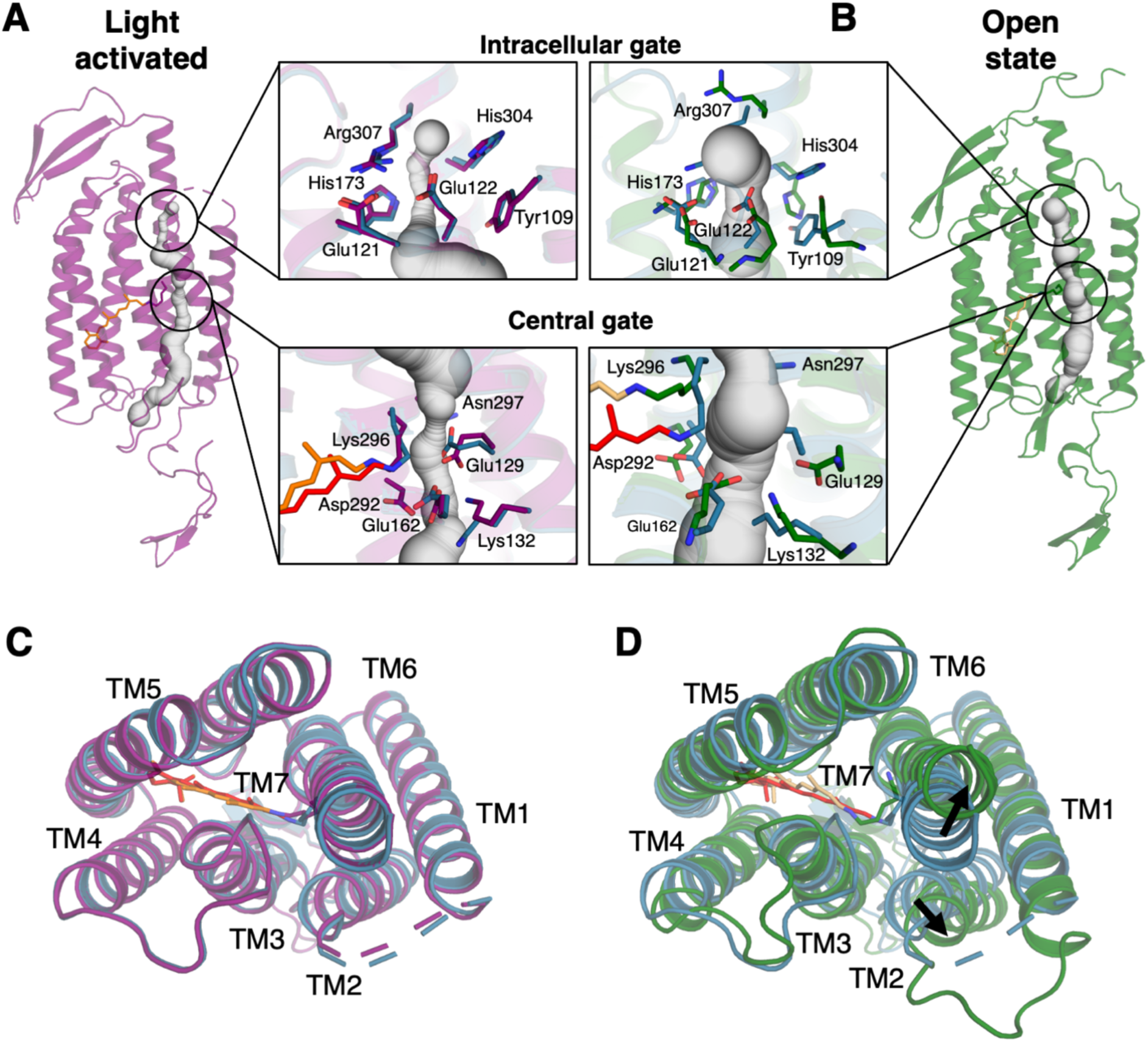
Opening the ion-conduction channel. (**A**) In the light-activated C1C2 structure (purple cartoon), conformational changes along the putative ion-translocation channel (grey) are predominantly located in the central gate (lower inset), leaving a bottleneck of 1.4 Å at the intracellular gate (upper inset). (**B**) Adjusting the protonation state based on spectroscopic characterizations of the open state (N_520_) leads to significant rearrangements in molecular dynamics simulations (right, green cartoon). The channel has further widened to a minimal width of 2.4 Å along the major bottlenecks sufficient for K^+^ ions translocation (compare Figure 3). The insets show a superposition of structures from the dark state (teal), light-activated state (purple), and open state (green), with key residues and the retinal chromophore depicted as sticks. (**C**) The overlay of the dark (teal) and light-activated (purple) C1C2 crystal structures indicates no large-scale rearrangements in the 7TM helical bundle. (**D**) In contrast the open state (green) is characterized by dominant shifts in TM2 and TM7 (black arrows) to open the cation conducting channel.

### Opening the cation conducting channel

To elucidate the conformational changes required to fully open the channel, we performed atomistic molecular dynamics (MD) simulations starting from the light-activated structure. In one set of simulations, we deprotonated the retinal Schiff base with Glu122, Glu129, Asp195, and Asp292 protonated to represent the M_390b_ state. The second set of simulations was biased towards the open N_520_ conformation, with a protonated retinal Schiff base and Asp292, but deprotonated Glu122, Asp129, and Asp195. As a further control, we performed the same simulations starting from the dark state structure. The different protonation states for these simulations were selected according to previous spectroscopic and electrophysiology experiments^12,14,28,42^ and are summarized in **Supplementary Table 2**. For each of these three distinct simulation setups, we embedded the channel into a palmitoyloleoyl phosphatidylcholine (POPC) lipid bilayer, surrounded by 600 mM KCl, under transmembrane voltages of 300-500 mV (one C1C2 channel under positive voltage, the other one negative (**Figure3D**)). Then we conducted five independent runs of 2 microseconds each. Relatively high transmembrane voltages compared to physiological conditions were used to accelerate channel opening and ion conduction within the simulated timeframe, a strategy extensively used in recent computational investigations of ion channels and membrane proteins^43–46^. Our MD simulations showed system stability throughout the simulation period, as evidenced by the backbone root mean squire deviation (RMSD) plots (**Supplementary Figure 3**), which plateaued and remained less than 3 Å.

In both the dark-state and M_390b_ protonation state simulations, the channel remained closed with no K^+^ permeation events (**Supplementary Figure 4+5**). Even though the central gate and extracellular vestibule in the light-activated structure were more open compared to the dark-state structure, the changes were not sufficient to allow K^+^ permeation (**Figure 2A**). Overall, the simulations thus supported our assignment of the light-activated structure to a non-cation-conducting early intermediate.

More significant conformational changes (root mean square deviation of backbone atoms in TM helices = 0.85-0.28 Å in the latter 1 µs of five 2 µs simulations) occurred in the set of simulations where we biased the structure from time-resolved serial crystallography towards the N_520_ intermediate by altering well-characterized protonation switches (**Figure 2B, Supplementary Table 2**). During these simulations, we observed substantial differences in the retinal binding pocket in both the central and intracellular gate (**Supplementary Figure 6**). Interestingly, both Trp262 and the interacting retinal chromophore, were considerably more dynamic in the open state simulations compared to the dark-state ones and the already significant conformational shift in the light-activated structure (**Supplementary Figure 7+8**).

These local changes are accompanied by a pronounced opening of the 7TM bundle compared to the dark and light-activated structures (**Figure 2 C, D**). The most significant changes were observed in the intracellular regions of TM1, TM2, TM6, and TM7, as well as the extracellular region of TM2, as evidenced by the per-residue backbone deviations calculated at different time frames of the MD simulations starting from the light-activated structure (**Supplementary Figure 9**). The conformational change in the opening of C1C2 notably differs from that reported in a recent study of the anion channelrhodopsin GtARC1^47^. Based on QM/MM and MD simulations, this study predicted a more localized conformational change around the retinal, leading to the opening of the constriction site in the ion permeation pathway. In contrast, the magnitude of changes in C1C2 are more akin to those observed during the formation of the open N-state structure of BR, where rearrangements in TM5, TM6 and TM7 initiate reloading of the proton pump^26^.

In C1C2 these changes are necessary for ion conduction and accordingly we observed 28 continuous K^+^ inward permeation events during the open state simulations (**Figure 3A**). This corresponds to a very low conductance of 1.2±1.1 pS, which falls within the range of experimentally estimated single-channel conductance^48,49^. Strong inward rectification was observed in the simulations, with very low outward conductance (**Supplementary Figure 10)**, which aligns well with the electrophysiology data^50^. Furthermore, our simulations revealed strict cation selectivity, with strong K^+^ binding but no Cl^-^ binding within the channel pore, likely due to the strongly electronegative environment (**Figure 3B, Supplementary Figure 11**). Moreover, we identified two neighboring stable K^+^ binding sites on the extracellular side of the channel pore, close to the central gate. At site 1, K^+^ is coordinated by Glu162, Thr289, Gln95, and Thr98, while at site 2, K^+^ is mainly coordinated by Glu162, Glu129, and Asp292 (**Figure 3C**). During conduction, K^+^ density within the pore was very low, with only one K^+^ ion conducted through the pore most of the time (**Supplementary Movie 1**). This conduction mechanism differs from those observed in most other cation channels, such as K^+^, Na^+^, and several non-selective cation channels^43,44,51,52^. These channels are multi-ion, single-file pores, where multiple ions permeate through this narrow region via a water-mediated or direct knock-on mechanism. In contrast, in Chlamydomonas the photocurrent is graded over four orders of magnitude in light intensity^53^ which requires channelrhodopsin to function as a one-ion pore, operating by simple mutual exclusion and a very low conductance.

**Figure 3:**
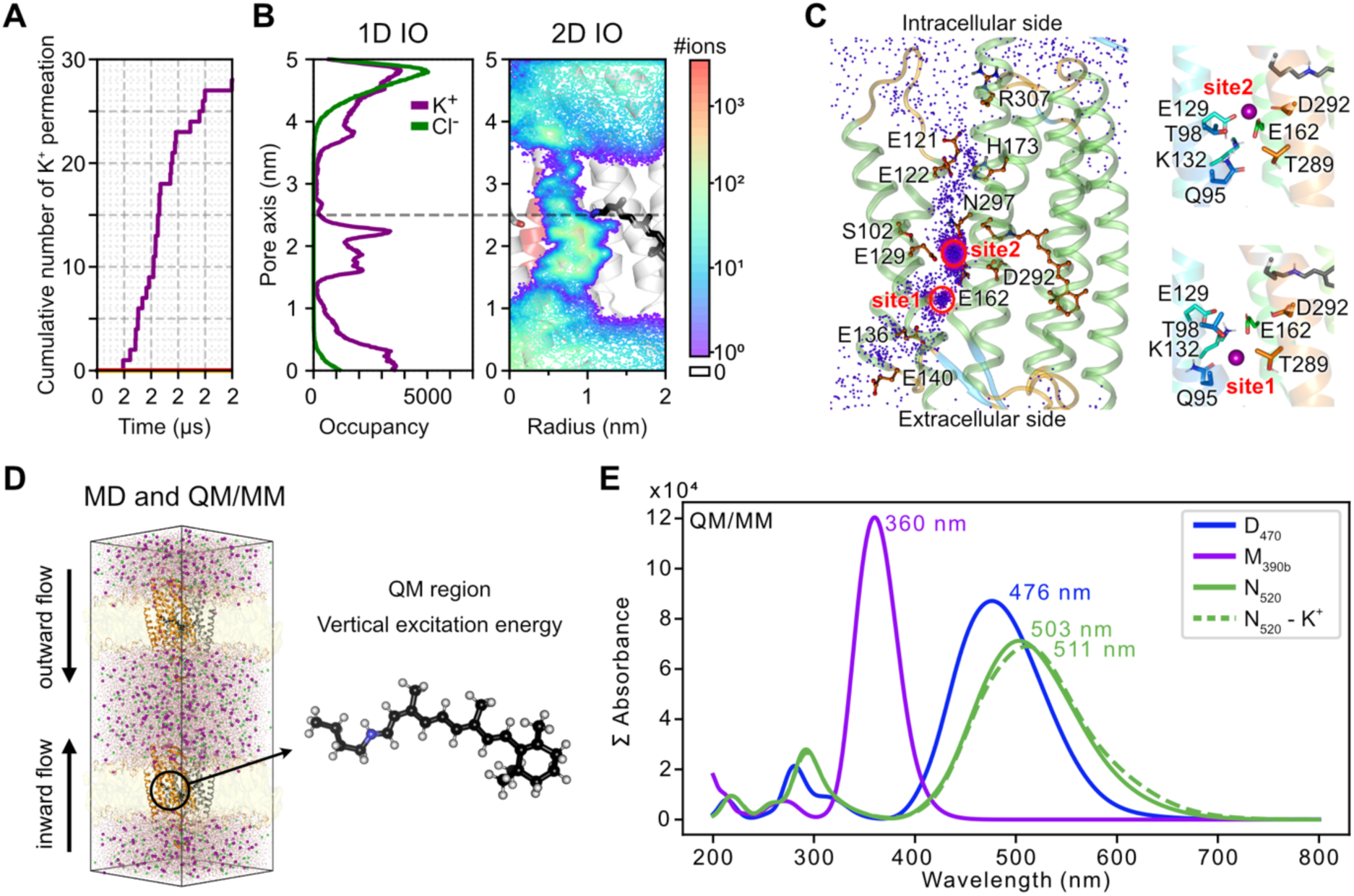
Potassium ion conduction through the open C1C2 pore. (**A**) Cumulative number of inward K^+^ permeation events passing through the C1C2 pore (orange dark, red light-activated, purple open state). (**B**). (Left) One-dimensional K^+^ and Cl^-^ ion occupancy along the pore axis derived from the open state simulations. (Right) 2D projection of ion occupancy within the pore cylinder with a radius of 2 nm and a height of 5 nm centered at the Cα of Ser102. The protonated retinal Schiff base and Ser102 are represented as stick models. (**C**) (left) Cumulative K^+^ ions passing through the C1C2 pore, derived from MD simulations, are depicted as purple spheres and mapped onto the end snapshot. (Right) Two major K^+^ binding sites revealed by MD. (**D**) The system setup for computational electrophysiology and hybrid quantum mechanics/molecular mechanics (QM/MM) simulations. Dimeric C1C2 is represented as gray and orange cartoon models, with POPC shown as yellow surface. Water molecules, K^+^ ions, Cl^-^ ions are depicted as red and white sticks, purple spheres, and green spheres, respectively. A charge imbalance of 6 *e* resulted in the membrane potential of ±389±27 mV across the two compartments (inner and outer). The QM region for the hybrid QM/MM simulations and vertical excitation energy calculations is depicted as a ball-and-stick model: The Cα and Cβ bond of retinal Schiff base was cut and capped with a hydrogen link bond. (**E**) Averaged UV-Vis spectra calculated from DFTB2 with dispersion correction. This excitation is on the ADC(2)/cc-pVDZ level of theory.

As a further step, we evaluated the different intermediates obtained from time-resolved serial crystallography and MD by predicting the vertical excitation energy using a hybrid quantum-mechanical molecular mechanics (QM/MM) approach (**Figure 3E**). The predicted maximum absorption of the light-activated state with deprotonated retinal was 360 nm slightly lower than experimentally observed difference of 390 nm but both are similarly blue-shifted compared to the dark-state. Furthermore, the MD-predicted open state showed a λ_max_ of 503 nm when no K^+^ is bound at site 2 and 511 nm when one K^+^ is bound at site 2. The difference between the open state and dark state is in good agreement with the experimentally characterized differences in the maximum absorption.

### Conclusions

In our study, we combined time-resolved serial crystallography with atomistic MD simulations to elucidate the major structural events during C1C2 channelrhodopsin activation. Within the crystal environment, C1C2 remained in a light-activated state that we assigned to an M_390_-like intermediate, where changes in the retinal binding pocket partially opened the central gate towards the extracellular vestibule.

While these structural changes widened the putative channel through the protein and may conduct H^+^, they were not sufficient to enable the flow of K^+^ in our simulations. Only when we set protonation switches in the central and intracellular gates, and DC pair to represent the N_520_ intermediate did we observe conformational changes that opened the intracellular gate, establishing a full channel for alkali cation but not anion conductance. The necessary structural changes can be illustrated by a morph between the light-activated structure and the simulated open state (**Supplementary Movie 2).** The mechanism, where initial light-induced changes partially open the channel but further protonation changes and larger helix arrangements are required for conductance (**Figure 4**), highlights the complex nature of channelrhodopsin activation.

**Figure 4:**
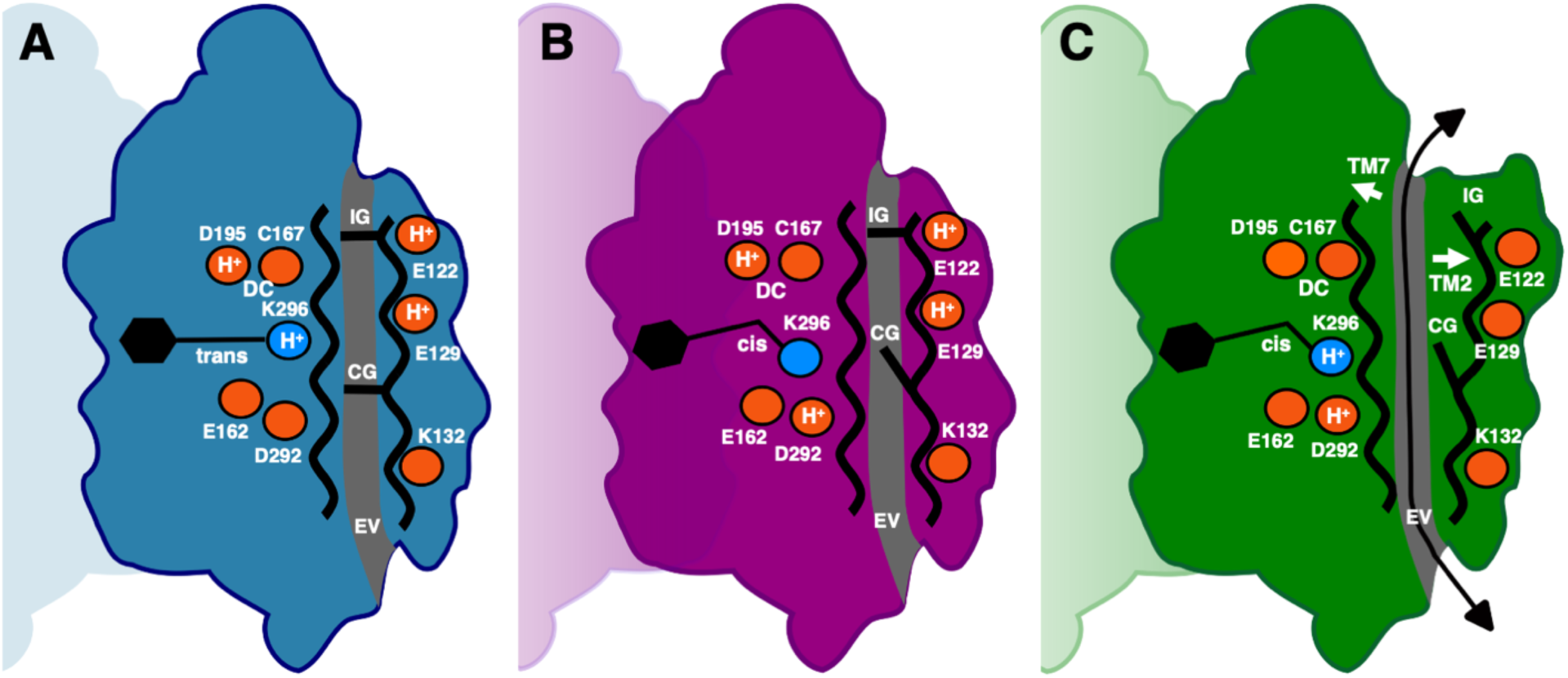
Schematic representation of the C1C2 opening mechanism. (**A**) In the dark state, (**B**) the M_390b_ intermediate and (**C**) the N_520_ intermediate. The retinal is shown in black and key residues discussed in the main text are shown as small rings. Protonation states (H^+^) used during the MD simulations are shown. The intracellular gate (IG), the central gate (CG) and the extracellular vestibule (EV) along the putative ion conduction channel (grey) are indicated. Opening of the channel is accompanied by rearrangements of the seven-transmembrane helical bundle and in particular of TM2 and TM7 (white arrows).

The general agreement between the simulated cation conductance and electrophysiology data, together with the absorption spectra calculated from our structural intermediates, suggests that this series of events closely resembles the structural changes upon channel opening. Further investigations could focus on capturing more intermediate states, on the conductance of H^+^ and Ca^2+^ and exploring the impact of different environmental conditions on channel activation. Extending this integrated structural approach to other channelrhodopsin variants could provide a broader understanding of the molecular principles underlying channel activation and how they could be exploited for designing optogenetic tools with tailored ion conducting properties.

## Materials and Methods

### Protein expression and purification

The human codon-optimized sequence of chimeric channelrhodopsin C1C2^4^, including a C-terminal tobacco etch virus (TEV) protease cleavage site followed by an octa-histidine tag, was cloned into the pFastBac1 vector (Invitrogen, Thermo Fisher Scientific, Waltham, MA, USA) using BamH1 and HindIII restriction sites. Protein expression in insect cells was carried out according to the Bac-to-Bac method, with bacmid preparation, transfection, virus production, and expression in Sf21 cells (Gibco, Thermo Fisher Scientific, Waltham, MA, USA), as described previously ^54^. After harvesting the cells all subsequent purification steps were performed at 4°C under red light. Cells were solubilized for 4h in buffer A (20 mM Tris-HCl pH 8.0, 100 mM NaCl) supplemented with 2% n-dodecyl-β-D-maltoside (DDM, Glycon Biochemicals GmbH, Luckenwalde, Germany), 0.4% Cholesterol hemisuccinate (CHS, Sigma-Aldrich, St. Louis, MO, USA), 5 µM all-trans retinal (Sigma-Aldrich) and complete protease inhibitor (Roche, Basel, Switzerland), and subsequently centrifuged at 40.000rpm for 30 minutes. The solubilized protein in the supernatant was purified by Ni^2+^ affinity chromatography (5ml HisTrap crude column (Cytiva, Marlborough, MA, USA)) followed by size-exclusion chromatography in buffer A with 0.05% DDM, 0.01% CHS (HiLoad 16/600 Superdex 200 pg (Cytiva)). Peak fractions with an absorption ratio of ∼ 2 at 280/475 nm were pooled, concentrated using an Amicon Ultra Centrifugal Filter with a molecular weight cutoff (MWCO) of 100 kDa (Merck Millipore, Burlington, MA, USA) and flash frozen in liquid nitrogen.

### Crystallization

We used a similar protocol for the crystallization of C1C2 as reported in^4^. In short, C1C2 was mixed with monoolein in a 2:3 protein to lipid ratio (w/w), in a 100 µL Hamilton syringe. The protein-LCP mixture was injected into crystallization buffer, containing 100 mM sodium citrate (pH 6.0), 30% PEG500DME, 100 mM MgCl_2_, 100 mM NaCl, and 100 mM (NH_4_)_2_SO_4_. Crystals were grown in 2-3 weeks in the dark at 20 °C. Crystal-laden LCP was pooled in a 500 µL Hamilton syringe with a small amount of crystallization buffer.

### Serial data collection

Dark and light-activated data were collected at the PXI-X06SA beamline at the Swiss Light Source with the setup described in^26^. To prepare samples for serial crystallography, the crystal-laden LCP was mixed with monoolein to reach the right consistency for extrusion. Next, the crystal-laden LCP was injected in the intersecting X-ray and laser diode paths by a high viscosity injector^55^ with a 50 µm diameter nozzle at a speed of 500 µm/s. Data was recorded on an EIGER 4M detector with a frame rate of 50 Hz. The X-ray beam had a size of 5x15 µm^2^ at an energy of 12.4 keV. A 445 nm laser diode was used for illumination of the crystals with a spot size of 80 x 120 µm, with a laser power of 4.9 mW/mm^2^, and an exposure time of about 100 ms. The time-resolved data was collected using a detector frame rate of 200 Hz using 200 ms cycle consisting of a 5 ms activation pulse from the laser followed by 39 bins of data.

### Serial data processing and refinement

Peak finding and indexing were performed with CrystFEL^56^, version 0.10.2, using the following settings: --indexing=xgandalf --peaks=peakfinder8 --threshold=10 --int-radius=4,5,7 --min-snr=3.5 –min-peaks=10 --min-pix-count=2 --min-res=80 –tolerance=5,5,5,3.0. Data was merged and scaled using partialator with model xsphere. The staraniso server was used for anisotropic correction of obtained data^57^. Isomorphous difference maps and extrapolated maps were calculated with Xtrapol8^58^, using k-weighting, and were used to aid refinement of the light structure. Structural refinements were done using Phenix^59^, version 1.20-4459, employing iterative cycles of manual adjustments made in Coot^60^. For final refinement of the light model the model refined against extrapolated data was combined with the dark model with an occupancy of 0.3 and 0.7 respectively and only B-factor and TLS refinement was carried out. All figures were created using the PyMOL Molecular Graphics System, Version 3.0 Schrödinger, LLC.

### Identifying channels in C1C2

Channels in the structures were identified using the Caver web tool^61^. Caver identified a pocket around the extracellular vestibule that acted as the starting point for the channel, with coordinates: x=3.7, y=20.2, and z=13. The settings for caver to identify the channel in the light-activated structure were: minimum probe radius = 0.7, shell depth = 4, shell radius = 3, clustering threshold = 3.5, maximal distance = 3, and desired radius = 5. We chose a minimum probe radius of 0.7 to showcase the bottleneck around the central gate as this is too small for alkaline cations to pass, as illustrated in **Figure 3**. For the simulated open structure, we changed the minimum probe radius to 0.8, the smallest size that allows alkaline cations to pass.

### Molecular dynamics (MD) simulations

We prepared systems for MD simulation of the dark-state (PDB entry 7C86) and our M-like structure (PDB entry 9GO2) of C1C2 using CHARMM-GUI^62^ with varied protonation states (**Supplementary Table 2**). For both the dark and open states, the missing region from Q112 to T117, as well as G329 and G330 was modeled using MODELLER 10.^63^. The dimeric C1C2 was embedded into 1-palmitoyl-2-oleoyl-sn-glycero-3-phosphocholine (POPC) and solvated with TIP3P water model^64^ and 600 mM KCl. Retinal force field parameters from the published works^65–67^ were used. CHARMM36m^68^ was employed for the simulations. The prepared systems (**Supplementary Table 3**) were energy minimized using the steepest descent method and equilibrated in an isothermal-isobaric (NPT) ensemble with a stepwise release of positional and dihedral angular restraints on the backbone, side-chains of the protein, and POPC (**Supplementary Table 4**). To maintain a constant temperature at 303 K and pressure at 1 bar during the equilibrations, we used the velocity-rescaling thermostat^69^ and Berendsen or Parrinello-Rahman barostats^70^. For the short-range van der Waals and Coulomb interactions, a cutoff of 1.2 nm was set. The particle mesh Ewald (PME) method^71^ was used for long-range electrostatic interactions. The LINCS algorithm^72^ was used to constrain the bonds between hydrogens and heavy-atoms. For the preparation of a system for MD-based computational electrophysiology (CompEL) simulations^73^ a copy of the equilibrated system was stacked along the channel pore axis, resulting in a double lipid bilayer system. This system was energy minimized, followed by production simulations for 2 μs with two charge imbalances (4 *e* and 6 *e*). The resulting transmembrane voltages of each simulation setup are summarized in **Supplementary Table 5**. The boundaries of the two compartments, inner and outer, were defined by the center of mass of each group of Cα atoms of the central gate residues (Ser102, Glu129, and Asn297) of the dimer. To maintain a constant charge imbalance across the membranes, ions crossing a cylinder centered at the defined group with radius of 3 nm and an upper and lower height of 1.5 nm and 1 nm, respectively, were counted. Upon detection of a crossed ion, an ion of the same species in one compartment was exchanged with water in the other compartment using the CompEL algorithm. These simulations were individually replicated five times. All MD simulations were conducted using GROMACS 2019.3 and 2019.6^74^.

### Analysis of the MD simulations

The plots of ion track (tracking the z-position of ions during their permeation), 1D and 2D ion occupancy were calculated using MDAnalysis 2.7^75^, Matplotlib 3.8.4^76^, Scipy 1.13^77^ and Numpy 1.26.1^78^. To track ions in the ion conduction pathway, ions within a virtual cylinder centered at the Cα of Ser102 with a radius of 2 nm and a height of 5 nm were considered. The protein was centered in the box, removing the periodic boundary effects, and then fitted to the backbone atoms in the initial snapshot for analysis. Molecular visualization was conducted using PyMOL 3.0.3 (http://www.pymol.org/pymol) and VMD 1.9.4^79^. For the deposition of the open state MD snapshot, we selected a representative structure from the third replica, which showed the largest conductance under a membrane potential of −389±27 mV. This selection was based on a conformational clustering using the *gmx cluster* module of GROMACS^74^, with a cutoff of 0.35 nm, resulting in 12 clusters. We chose the representative structure from the first cluster.

### Hybrid quantum mechanics / molecular mechanics (QM/MM) simulations

For the dark, light-activated, and open state, one trajectory of the MD runs was processed into 100 equidistantly chosen snapshots. For the light-activated state, five input structures for the QM/MM runs were selected from these snapshots. As the channel opening is of predominant interest, we processed more trajectories within the dark and open state characterization. For the dark state, eight trajectories were processed, while for the open state, the snapshots were divided into those with and without a potassium ion in close vicinity to the retinal chromophore (potassium ions were considered close, if they were found in a 6.5 Å distance of retinal Schiff base, Glu162, and Asp292). Again, eight structures of each subset were included in the QM/MM runs. The selected snapshots were chosen from the entire shift range of the optimized structures (see below).

All QM/MM simulations were performed using the AMBER suit of programs, version 20 or newer^80^. Throughout the hybrid QM/MM simulations, the QM region was treated either by the DFTB2 or DFTB3 tight binding model as implemented internally in AMBER^81–83^, using the MIO-L and 3OB parameter sets, respectively^84–86^. For DFTB2 runs, the implemented dispersion model was employed (DFTB2+D)^87^.

The QM region of the QM/MM runs included the retinal Schiff base, capped with hydrogen at the Cα atoms and embedded in a point charge environment, i.e. electrostatic embedding was used. All non-QM atoms were described with the CHARMM36m all atom force-field^68^ in all QM/MM simulations.

QM/MM runs are based on MD equilibrated snapshot structures, followed by QM/MM minimization with a short pre-minimization (250 steps; steepest decent algorithm) using constraints on the protein and lipids, followed by 250 steps of steepest descent and additionally 750 steps using the conjugate gradient method without any constraints on DFTB3 level of theory. The 100 ps equilibration was separated into a 50 ps heating step (100 K to 303 K) followed by 50 ps unconstrained equilibration. For each input structure, a production of 100 ps was performed. Equilibration and production runs were recorded predominantly on DFTB2+D level of theory. As a comparison and method validation for a subset of the data, equilibration and production was also performed on DFTB3 level of theory (**Supplementary Figure 12**). Since both computations lead to qualitatively similar results, we finally decided on DFTB2+D in order to benefit from the implemented dispersion correction.

Spectra were generated from equidistantly spaced snapshots (every 1 ps) of the production runs. In all cases, the retinal chromophore was extracted as the QM region, keeping the protein environment as a point charge embedding in the QM calculations. Spectra were calculated on RI-ADC(2)/cc-pVDZ^88^ level of theory with the Turbomole programsuit version 7.7 or newer (TURBOMOLE V7.7 2022, a development of University of Karlsruhe and Forschungszentrum Karlsruhe GmbH, 1989-2007, TURBOMOLE GmbH, since 2007.). The convergence of the self-consistent field (SCF) computations was set to 10^-8^, additionally ensuring converged results by enforcing changes of the density matrix to be smaller than 10^-7^. For SCF and ADC(2) excitation computations, the resolution of the identity approximation was used. Except for rare cases where convergence issues appeared (here the number of recorded excitations was throughout reduced to 8; 11 appearances in all calculations), the 10 lowest excited states were recorded. Spectra were computed as an average of all excitations and intensities were calculated from the oscillator strength of the individual excitations.

## Acknowledgements

We are grateful for the excellent support from the PSI Crystallization Facility and the Macromolecular Crystallography group throughout the optimization and crystal testing process. Demet Kekilli, Hannah Glover, Maximillian Wranik, Antonia Furrer and Daniel James assisted during X-ray beamtimes. We acknowledge Dr. Johannes Vierock from the Charité – Berlin University Medicine and Dr. Tillmann Utesch from FMP Berlin for the helpful discussion, as well as Dr. Johann Biedermann from FMP Berlin for the analysis of MD data. The authors gratefully acknowledge the computing time made available to them on the high-performance computer “Lise” at the NHR Center NHR@ZIB. This project was funded by the following agencies: the Swiss National Science Foundation under projects grants 310030_197674 (to T.W.) and 310030_207462 (J.S.); and Swiss Innovation Agency Innosuisse grant 42711.1 IP-LS (to J.S.). DFG under Germany’s Excellence Strategy EXC 2008–390540038 – UniSysCat and CRC1078 ‘Protonation Dynamics in Protein Function’ (to H.S.). The European Research Council (ERC)-2020-Synergy grant (“SOL” 951644, P.H.). P.H. is Hertie Professor and supported by the Hertie Foundation.

## Author contributions

Conceptualization: P.H., H.S. and J.S.

Protein Production: M.B., Ch.S. and P.H.

Crystallization: P.S. and S.B.

X-ray data collection: S.B., P.S., T.W. F.D. and J.S.

X-ray data processing: M.M., S.B. and T.W.

MD simulations: S.H and H.S.

QM/MM calculations: S.H., Ca. S., J.C., I.S. and H.S.

Writing - original draft: M.M., S.H., H.S. and J.S.

Writing - review and editing: All authors could read and comment on the manuscript.

## Competing interests

The authors declare no competing interests.

## Data and materials availability

Coordinates and structure factors have been deposited in the PDB database under accession codes 9GO1(for the C1C2 dark state) and 9GO2 (for the light-activated state). Data relevant to molecular dynamic simulations has been deposited in Zenodo linked under https://tinyurl.com/82uttvkv.

---

**Supplementary Figure 1:**
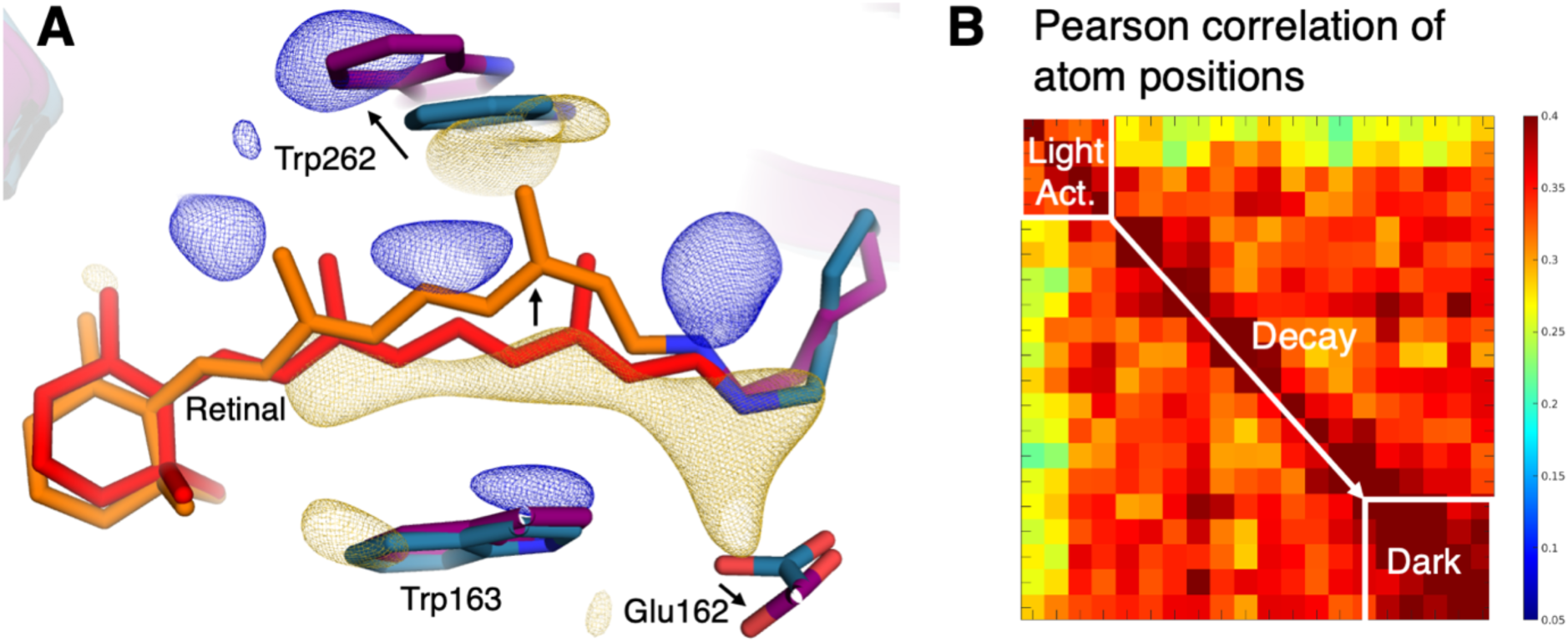
Difference electron density map and Pearson correlation of atom positions. (**A**) View of changes within the retinal binding pocket. Movements of the retinal chromophore (red dark state, orange light-activated state) and nearby residues (dark in teal, light-activated in purple) are indicated by arrows. The experimental difference electron density map (Fo(light)-Fo(dark), gold negative, blue positive, contoured at 3.5 sigma) from the serial crystallographic experiment is shown. (**B**) Additional time-resolved serial crystallographic data was collected as described in^26^ and binned into 10 ms intervals. Structural changes can be followed by the rmsd correlation of all atoms, including the retinal, in each binned structure. The results identify two major states at about 5-40 ms and 40-200 ms corresponding to the formation and decay of the light-activated structure.

**Supplementary Figure 2:**
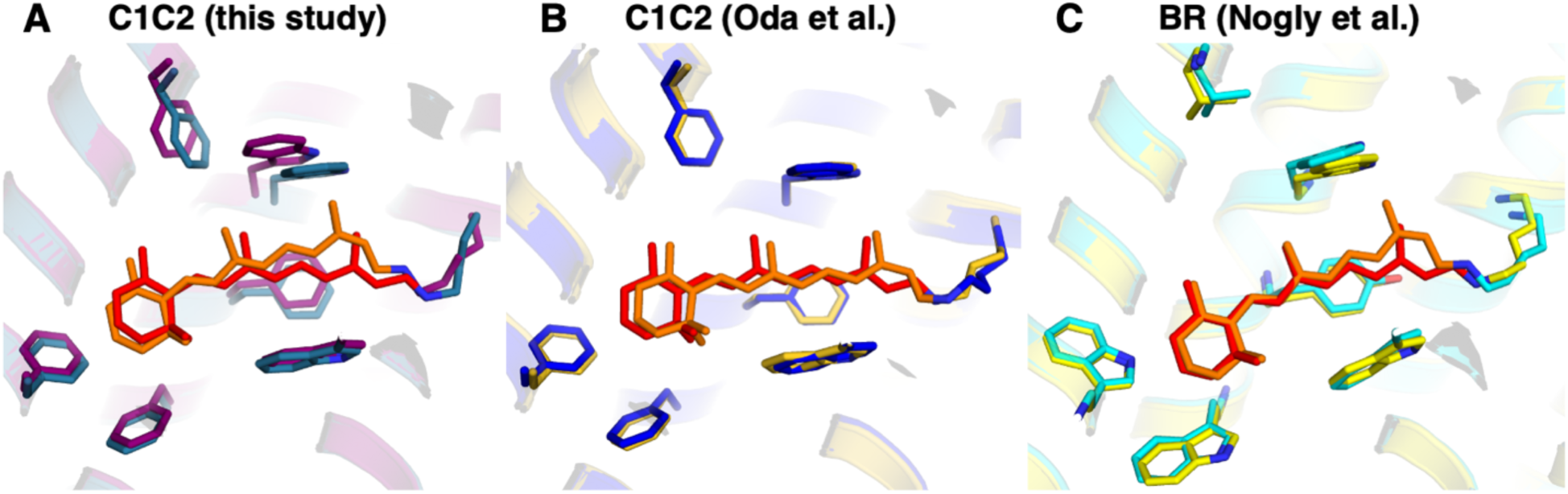
Comparison of light-activated rhodopsin structures. (**A**) Structures of C1C2 channelrhodopsin from this study (red dark, orange light-activated for about 100 ms). (**B**) Structures of C1C2 taken from^21^ (red dark, orange 4 ms after light-activation). (**C**) Structures of the BR taken from^32^ (red dark, orange 8.33 ms after light-activation). It is interesting to note how similar our early deprotonated structure of C1C2 is to the M-intermediate structure of BR. In contrast our structure is markedly different to the C1C2 structures reported by Oda et al., who refined their data against a 13-cis,15-syn retinylidene conformation and very limited changes in the retinal binding pocket.

**Supplementary Figure 3:**
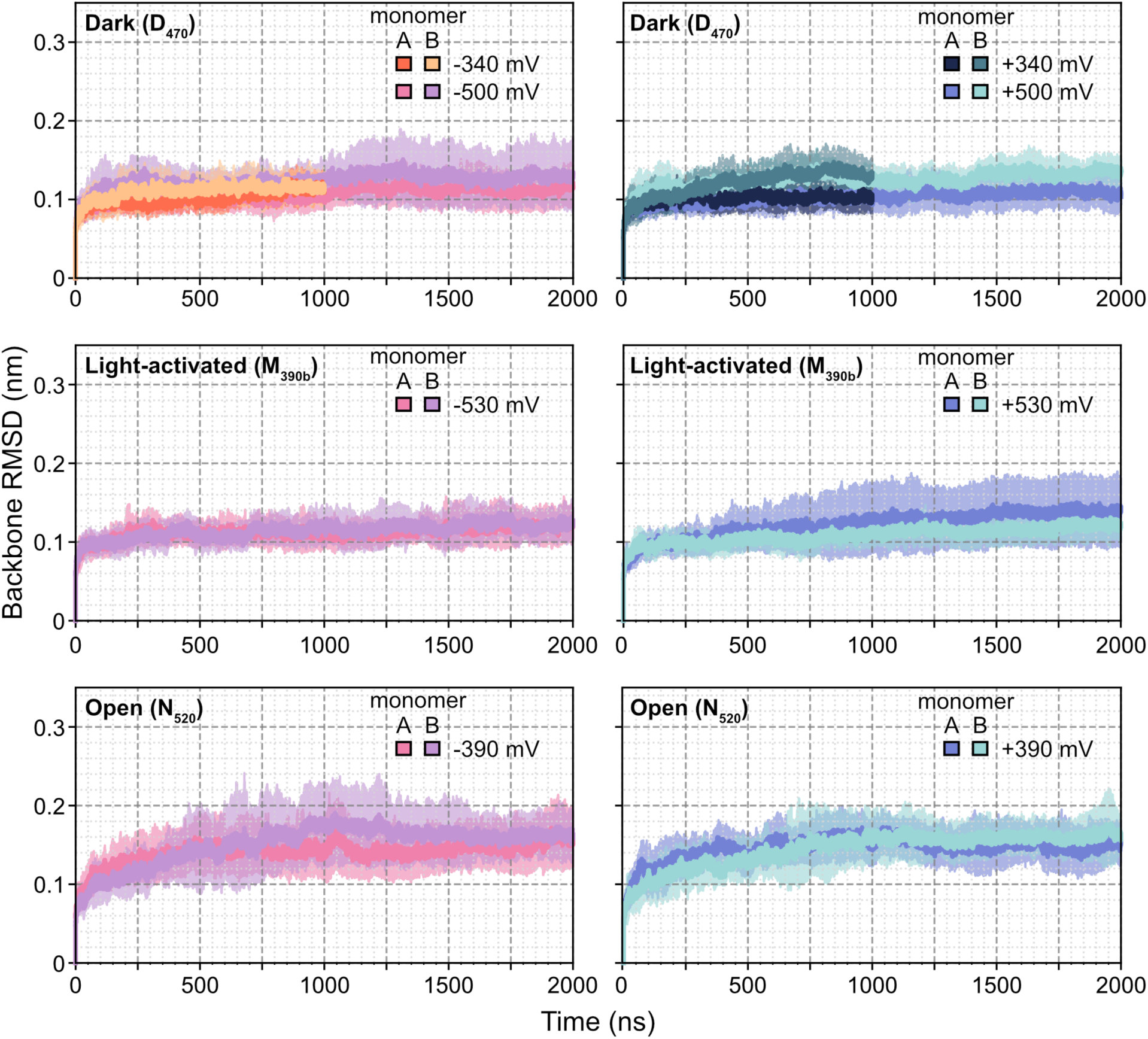
Root-mean-square-deviations (RMSDs) during molecular dynamic simulations. The average RMSDs of backbone of each monomer in the dimeric C1C2 in the dark state, light-activated, and open state are shown as solid lines, while the standard deviations are shown as shaded area. For these RMSD calculations, residues 85 to 308 were considered, excluding the flexible extracellular amino and intracellular carboxyl termini. The simulations were conducted for 2 µs, except for the dark state at ±340 mV, which was run for 1 μs. Each simulation was replicated five times at 303 K and with a 600 mM KCl concentration.

**Supplementary Figure 4:**
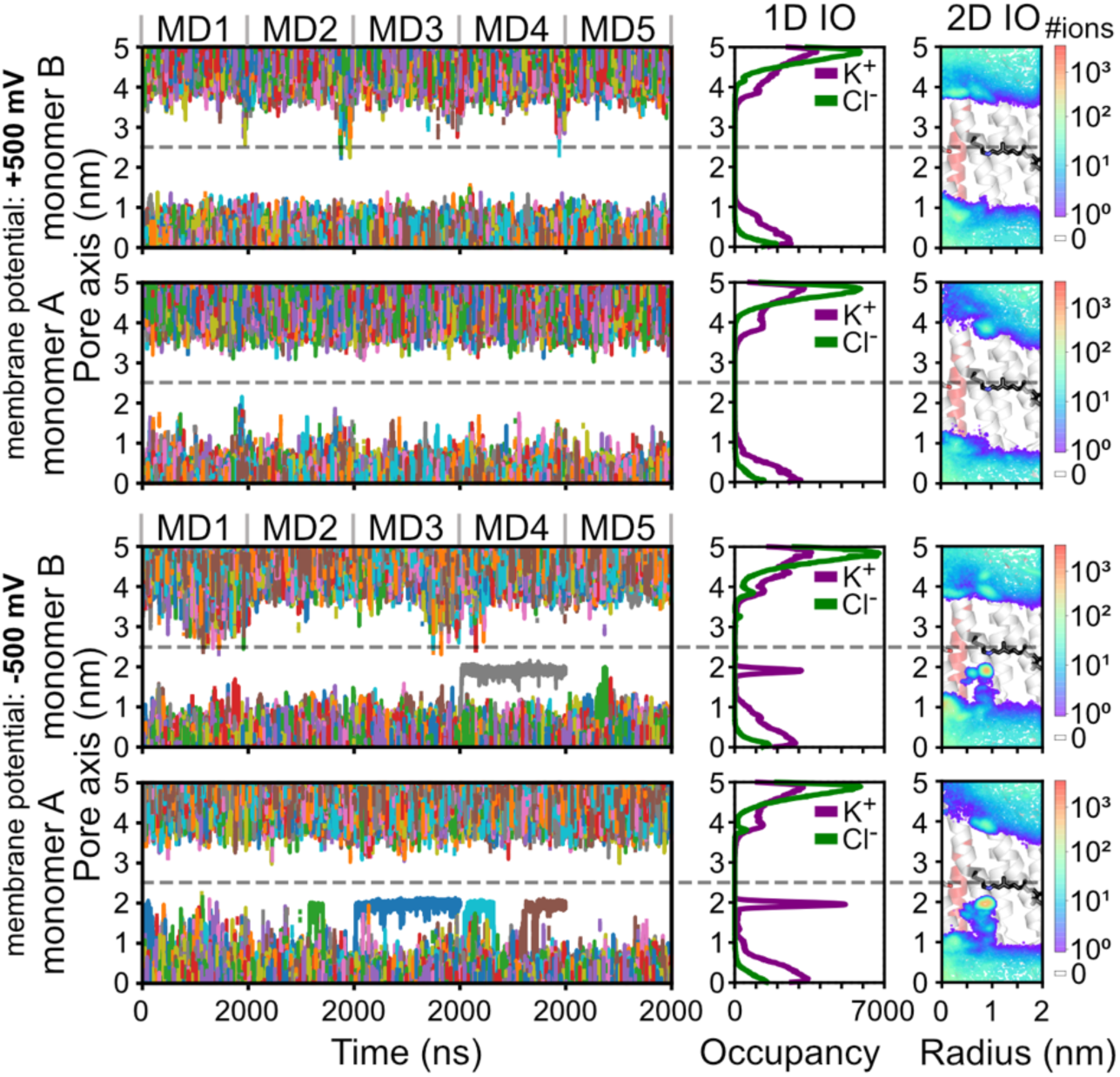
Cation permeation in the dark-state of C1C2. (**left**) Traces of K^+^ in the pore of C1C2 with the central gate highlighted as a dashed grey line. (**middle**) Cumulative 1D ion occupancy along the pore axis. (**right**) 2D K^+^ density within the pore region, mapped onto the closed-state C1C2, with protonated retinal Schiff base and Ser102 depicted as stick models. The 2-μs simulations were replicated five times at 303 K with a 600 mM KCl concentration. No ion permeation events were observed in these simulations.

**Supplementary Figure 5:**
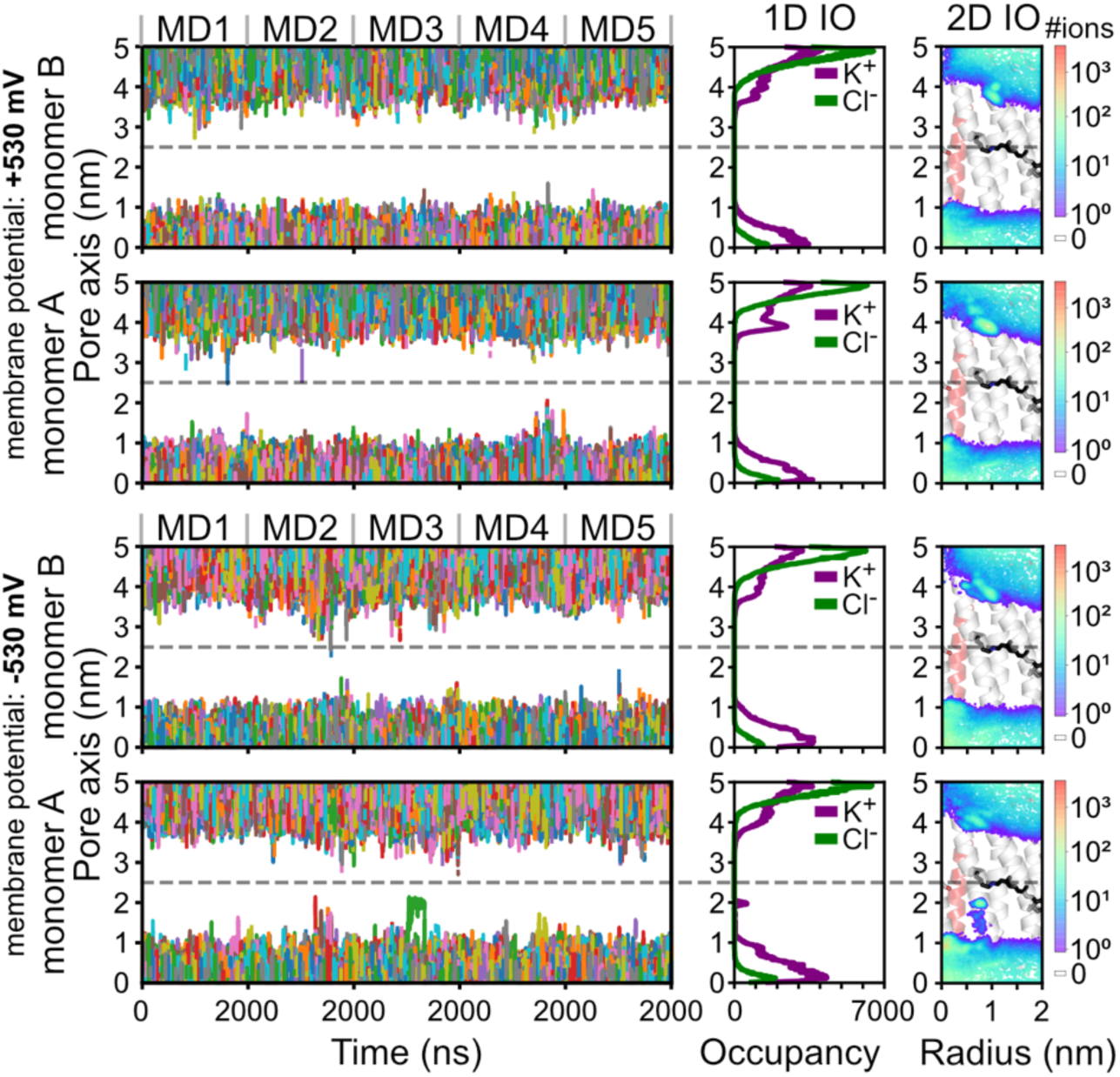
Cation conduction through the light-activated C1C2 structure. (**left**) Traces of K^+^ in the pore of C1C2 with the central gate highlighted as a dashed grey line. (**middle**) Cumulative 1D ion occupancy along the pore axis. (**right**) 2D K^+^ density within the pore region, mapped onto the closed-state C1C2, with protonated retinal Schiff base and Ser102 depicted as stick models. The 2-μs simulations were replicated five times at 303 K with a 600 mM KCl concentration. No ion permeation events were observed in these simulations.

**Supplementary Figure 6:**
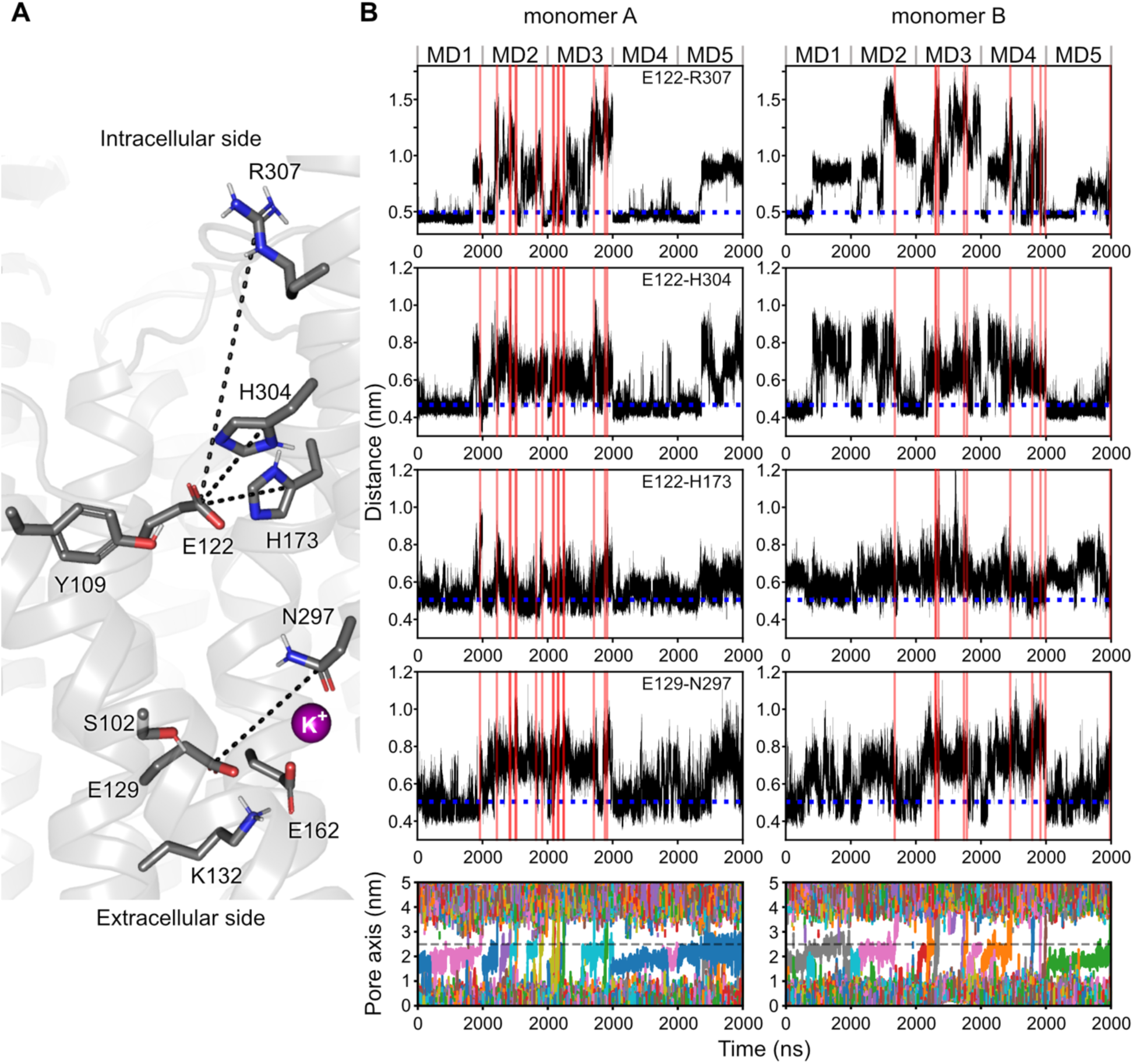
Time evolution of residue distances during light-gating, and K^+^ conduction events observed during the open state simulations. (**A**) Key residues considered for residue-pair distance measurements are depicted as stick models, and their calculated minimum distances are represented as dotted lines in the end snapshot of a 2-μs simulation of the C1C2 open state. (**B**) (from the first to the fourth rows) Time evolution of atomic distances of the key residues in the intracellular gate and central gate, and (in the fifth row) 1D ion tracking of K^+^ within each pore with a radius of 2 nm and height of 5 nm, centered at each Cα of Ser102 in monomer A and B. (from the first to the fourth rows) The red lines indicate the corresponding inward K^+^ permeation events passing through the central gate. The blue dotted lines represent the calculated minimal distances in our x-ray structure of M_390_. The 2-μs simulations were replicated five times at 303 K with a 600 mM KCl concentration under a membrane potential of −389±27 mV.

**Supplementary Figure 7:**
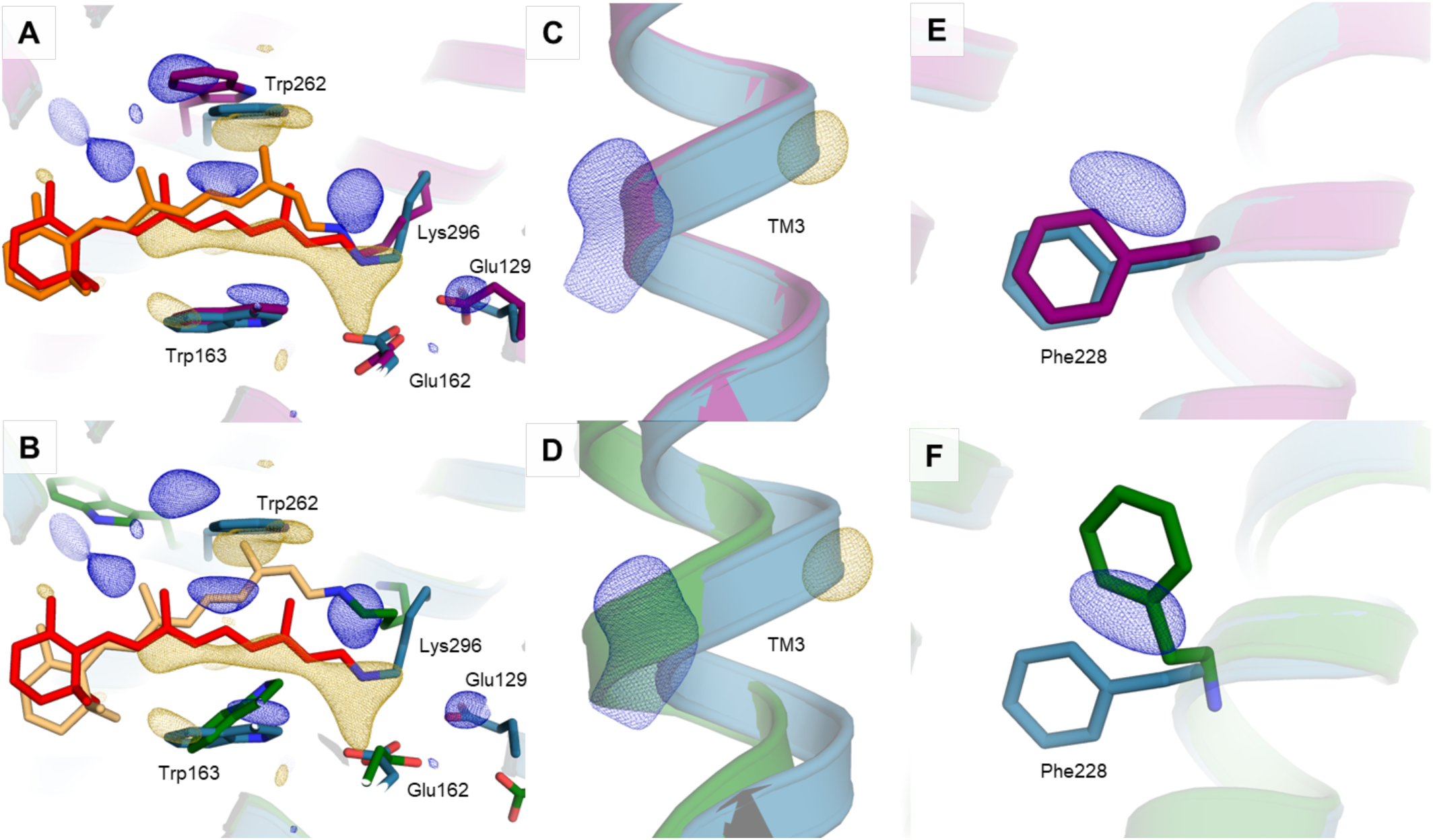
Contributions of open state to light-activated data. (**A**) The refined structures (teal/red for dark and purple/orange for light-activated) explain the difference electron density map ((Fo(light)-Fo(dark), gold negative, blue positive, contoured at 3.5 sigma) well. (**B**) Even though the refined light-activated structure is the dominating state, an overlay of the difference electron density map with the simulated open state (green/lightorange) suggests that both intermediates contribute and the open state can occur *in crystallo*. Further evidence can be found in the transmembrane helix 3 (**C** and **D**) and on Phe228 (**E** and **F**). These observations provide a cross-validation for the molecular dynamic simulations and open the possibility to resolve both states by time-resolved structural biology.

**Supplementary Figure 8:**
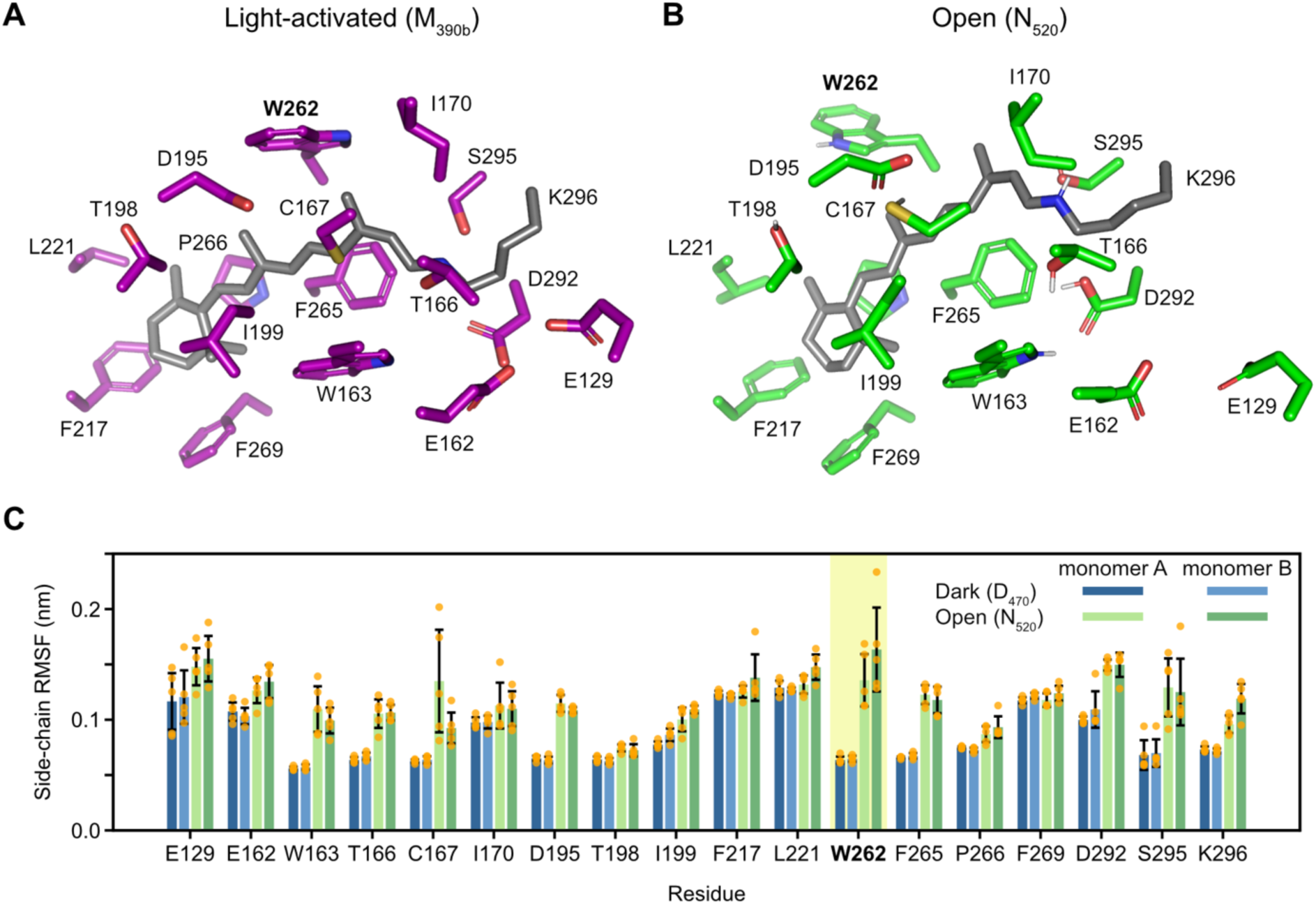
Side-chain dynamics of the retinal binding site in C1C2. (**A**) Residues in the retinal binding site within 4.5 Å of the retinal moiety in the simulated M_390b_ state were selected for the calculation of root mean square fluctuation (RMSF) as shown in C. (**B**) The corresponding residues in the open state. (**C**) Residue-wise RMSF of the side-chain in the retinal binding site was calculated for each monomer in both the dark and open states. The 2-μs simulations were replicated five times at 303 K with a 600 mM KCl concentration under a membrane potential of −503±30 mV for the dark state and −389±27 mV for the open state.

**Supplementary Figure 9:**
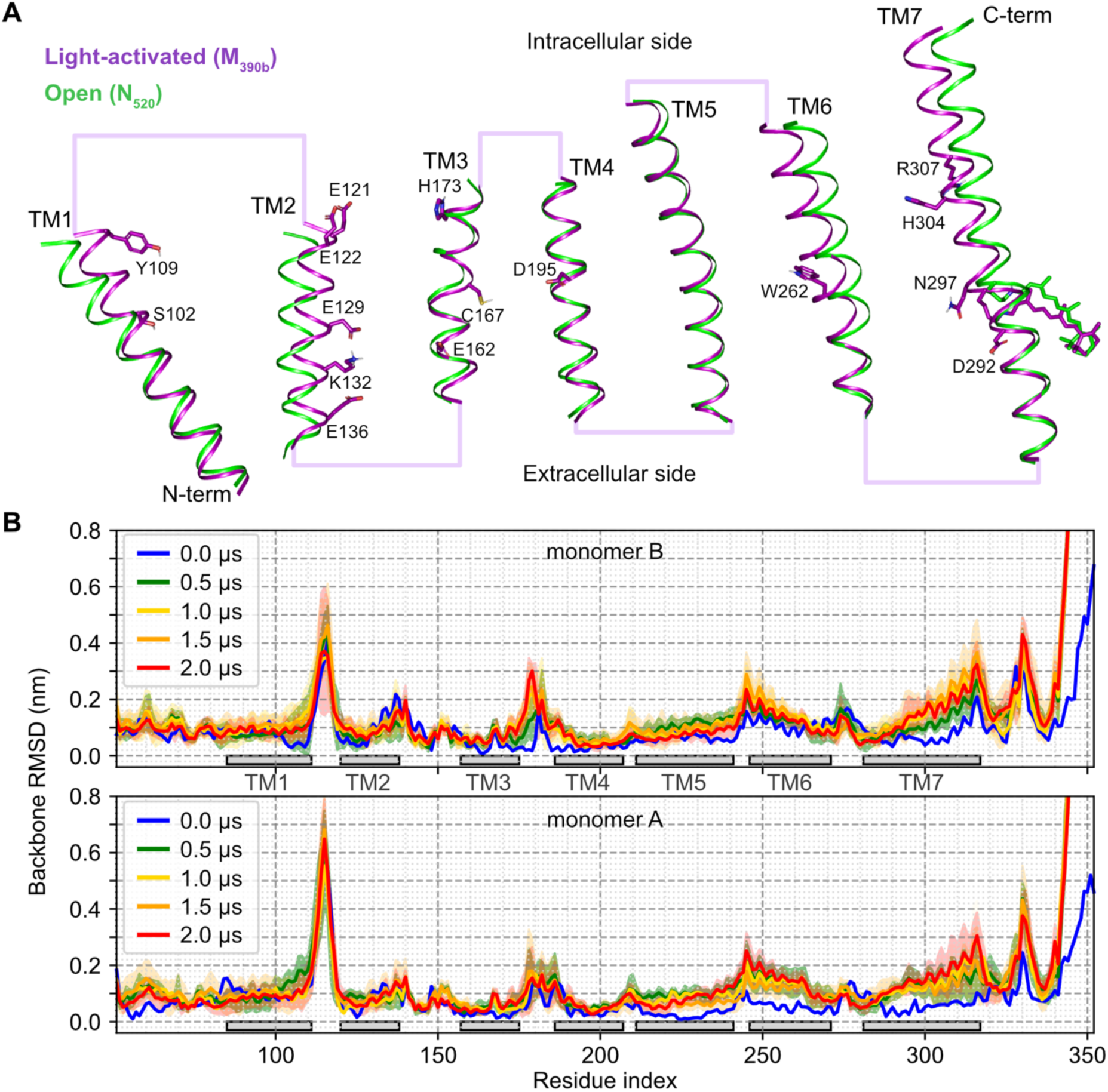
Conformational transition from the light-activated state to the open state. (**A**) Cartoon representations of the transmembrane helices of the light-activated structure (purple, PDB entry 9GO2) and the end snapshot of an open state simulation replica at 2 μs. (**B**) Per-residue backbone root-mean-square deviations (RMSDs) of each monomer in the open state during production runs at 0, 0.5, 1, 1.5, and 2 μs. The reference structure for the RMSD calculation was set to the backbone of the light-activated structure. The 2-μs simulations of the open state were replicated five times at 303 K with a 600 mM KCl concentration under a membrane potential of −389±27 mV. The solid line and shading represent the mean and standard deviation of the RMSDs.

**Supplementary Figure 10:**
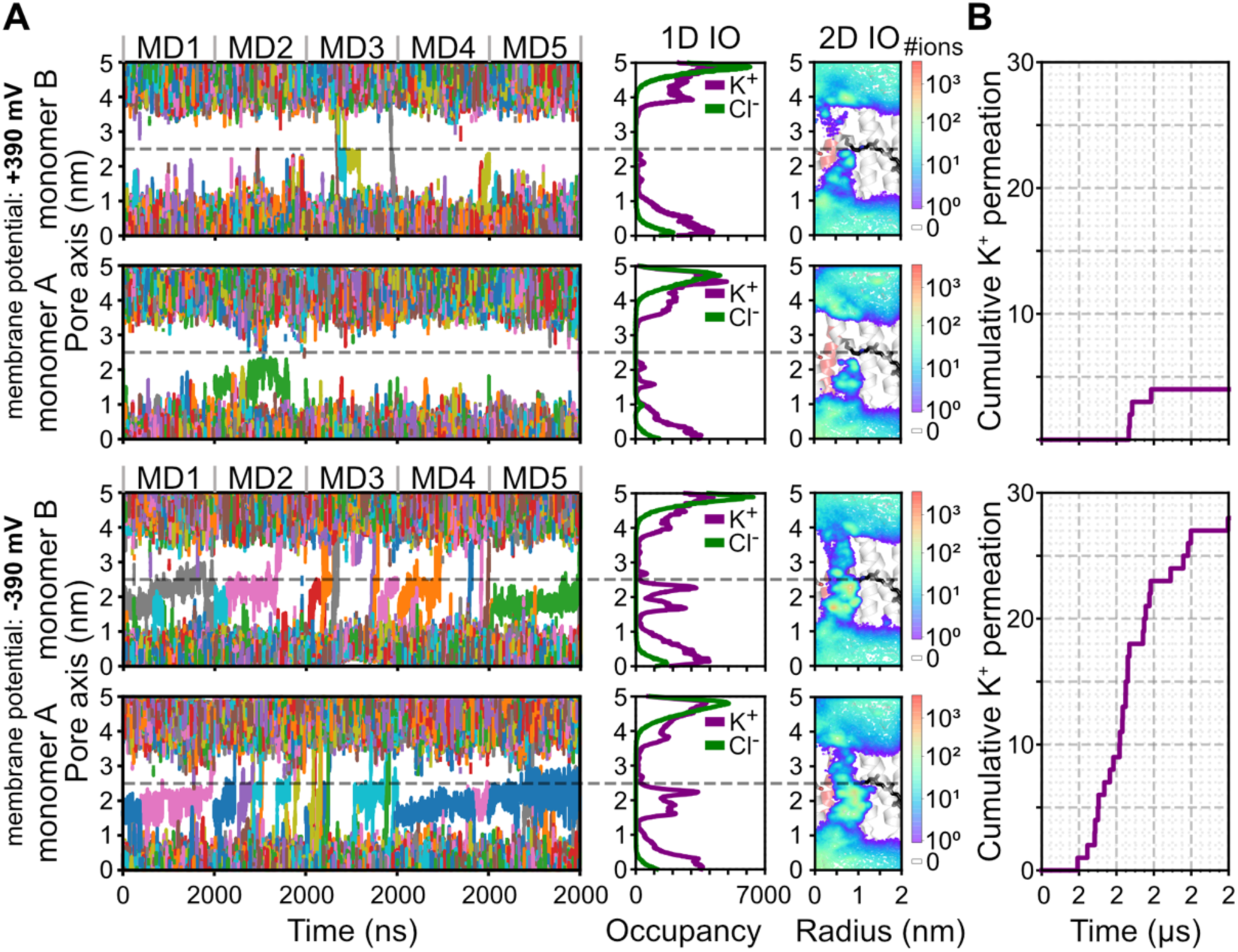
Cation conduction through the open state of C1C2. (**A**) (**left**) Traces of K^+^ in the pore of C1C2 with the central gate highlighted as a dashed grey line. (**middle**) Cumulative 1D ion occupancy along the pore axis. (**right**) 2D K^+^ density within the pore region, mapped onto the open state C1C2, with protonated retinal Schiff base and Ser102 depicted as stick models. (**B**) Cumulative number of outward and inward K^+^ permeation events passing through the central gate of C1C2 pore is shown in the upper and lower panels, respectively. (**A, B**) The 2-μs simulations were replicated five times at 303 K with a 600 mM KCl concentration.

**Supplementary Figure 11:**
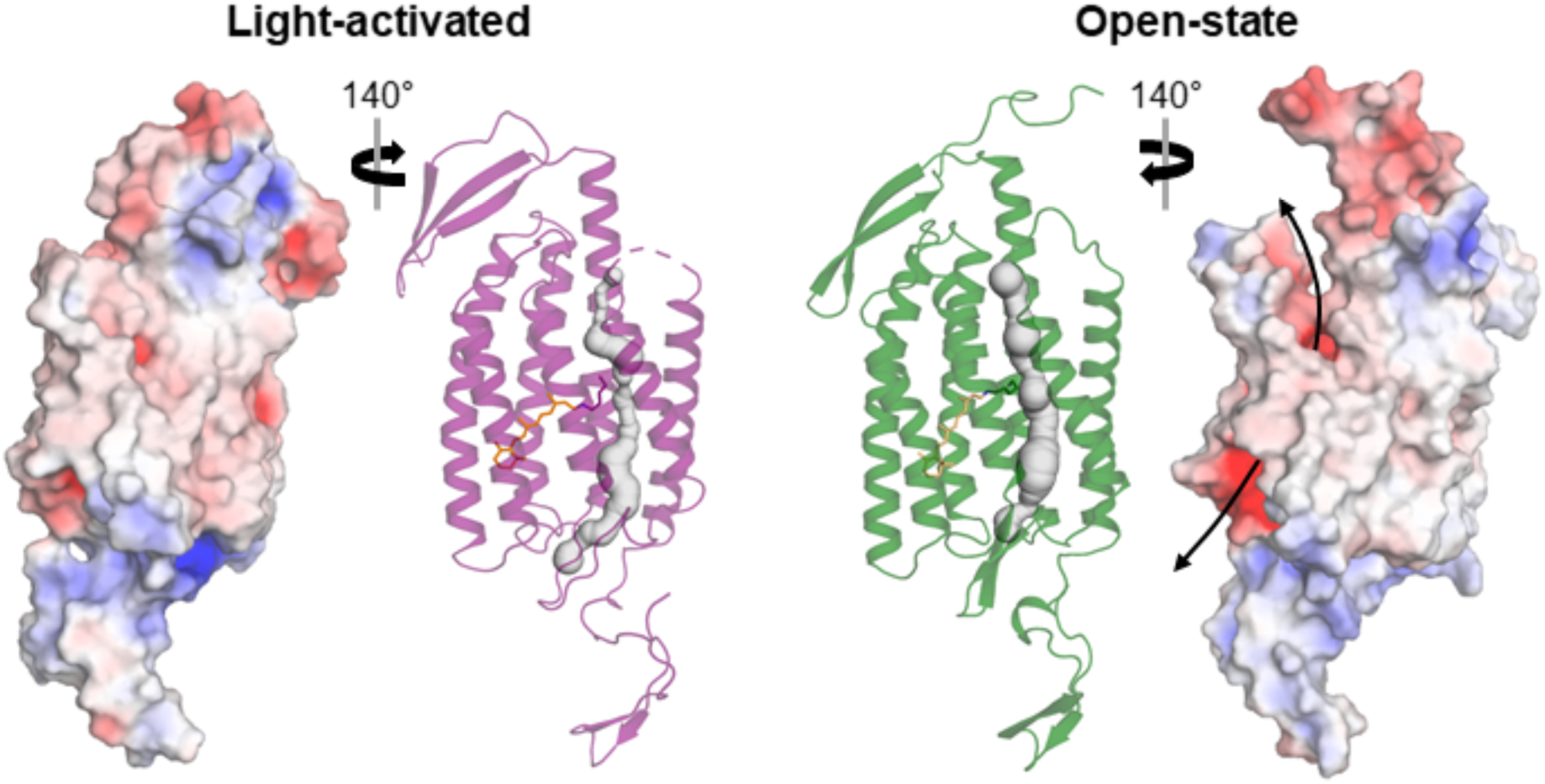
Comparison of the charge distribution in the light-activated and open states of C1C2. The light-activated C1C2 structure (purple cartoon) and its electrostatic potential map (**left**) are compared with the representative open-state structure obtained by clustering from the MD simulation under a membrane potential of −390 mV and its electrostatic potential map (**right**), along with their putative ion-translocation channels (grey). In the electrostatic potential maps, positively and negatively charged regions are colored blue and red, respectively. The cation translocation pathway, indicated by a black arrow, is lined by negative charge supporting the selective flow of cations.

**Supplementary Figure 12:**
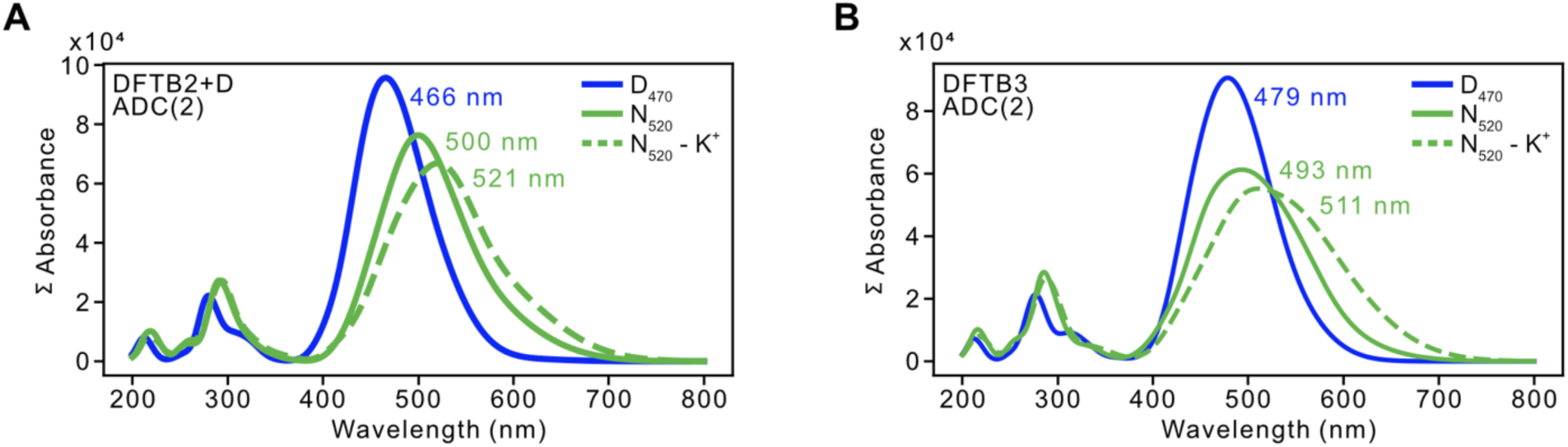
Averaged vertical excitation energies of the dark and open (with and without K^+^ in close vicinity of the retinal chromophore) states on RI-ADC(2)/cc-pVDZ level of theory, based on three QM/MM simulations runs for each state on (**A**) DFTB2+D and (**B**) DFTB3 level of theory.

**Supplementary Movie 1: Potassium ion translocation via the open state.** The MD trajectory from 850 ns to 875 ns of monomer A of the open state of C1C2 is shown. K^+^, Cl^-^, and water molecules are represented by purple, green, and cyan spheres, respectively. The key residues along the K^+^ translocation pathway - (from the bottom to the top) Glu140, Glu136, Lys132, Glu162, Glu129, Asp292, Ser102, Asn297, Glu122, Glu121, His173, His304, and Arg307 - are depicted as stick models. For clarity, monomer B and POPC lipids are not shown, and only the K^+^, Cl^-^, and water molecules within 5 Å of the monomer A are displayed.

**Supplementary Movie 2: Structural changes upon channel opening.** This morph between the dark structure, the light-activated structure and the MD simulation using N_520_ protonation states illustrates the structural changes in the retinal binding pocket, the central gate and the intracellular gate upon channel formation.

**Supplementary Table 1:** Crystallographic data and refinement statistics.

| Dataset | Dark | Light-activated |
| --- | --- | --- |
| **Data Collection** |  |  |
| Resolution (Å) | C222<sub>1</sub> | C222<sub>1</sub> |
| a, b, c (Å) | 61.34, 141.70, 94.40 | 61.34, 141.70, 94.40 |
| $\alpha$, $\beta$, $\gamma$ (°) | 90, 90, 90 | 90, 90, 90 |
| **Overall Statistics (High-Resolution Statistics)** |  |  |
| Resolution (Å) | 94.40-2.60 (2.70-2.60) | 94.40-2.70 (2.80-2.70) |
| Indexed patterns | 55249 | 43646 |
| Indexing rate (%) | 6.33 | 5.1 |
| No. Reflections | 13062 (1288) | 11685 (1150) |
| Completeness (%) | 100 (100) | 100 (100) |
| Multiplicity | 344.3 (139.2) | 215.40 (33.2) |
| Rsplit (%) | 7.93 (272.12) | 9.4 (359.25) |
| CC* | 0.999 (0.810) | 0.999 (0.796) |
| $\langle I/\sigma(I) \rangle$ | 7.39 (0.32) | 6.37 (0.25) |
| **Anisotropy Corrected Statistics (High-Resolution Statistics)** |  |  |
| Resolution (Å) | 70.85-2.60 (2.69-2.60) | 70.85-2.70 (2.80-2.70) |
| No. Reflections | 8838 (1288) | 7315 (72) |
| Completeness (%) | 67.7 (10.3) | 62.6 (6.3) |
| Multiplicity | 415.94 (167.4) | 348.47 (161.5) |
| Rsplit (%) | 6.08 (87.08) | 6.88 (84.77) |
| CC* | 0.999 (0.849) | 0.999 (0.75) |
| $\langle I/\sigma(I) \rangle$ | 10.7 (1.05) | 9.98 (1.16) |
| **Refinement** |  |  |
| Resolution Range (Å) | 70.85-2.59 | 70.85-2.70 |
| No. Reflections | 8,836 | 7312 |
| R<sub>work</sub> / R<sub>free</sub> | 0.2171 / 0.2696 | 0.2399 / 0.2682 |
| **No. Atoms** |  |  |
| Protein | 2332 | 2327 |
| Other | 126 | 101 |
| Water | 13 | 4 |
| **B Factors** |  |  |
| Protein | 54.6 | 42.8 |
| Other | 72.9 | 51.5 |
| Water | 40.8 | 31.9 |
| **R.m.s. Deviations** |  |  |
| Bond Lengths (Å) | 0.005 | 0.004 |
| Bond Angles (°) | 0.919 | 0.810 |
| **Ramachandran** |  |  |
| Favored (%) | 97.89 | 97.89 |
| Allowed (%) | 2.11 | 2.11 |
| Outliers (%) | 0.00 | 0.00 |
| PDB ID | 9G01 | 9G02 |

**Supplementary Table 2:**
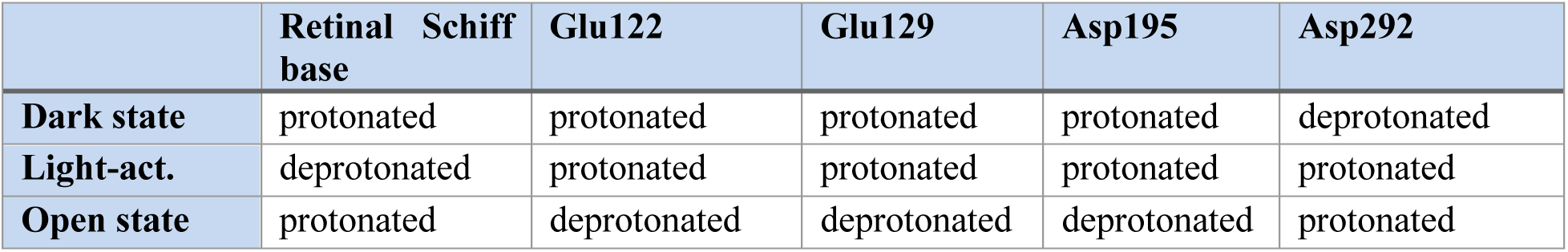
Protonation states used for MD simulations. . Protonations for the dark-state, the early light-activated intermediate, and the open state were selected according to previous spectroscopic and electrophysiology experiments^12,14,28,42^.

|  | Retinal Schiff base | Glu122 | Glu129 | Asp195 | Asp292 |
| --- | --- | --- | --- | --- | --- |
| **Dark state** | protonated | protonated | protonated | protonated | deprotonated |
| **Light-act.** | deprotonated | protonated | protonated | protonated | protonated |
| **Open state** | protonated | deprotonated | deprotonated | deprotonated | protonated |

**Supplementary Table 3:** Details of the CompEL simulation system. The simulations were conducted at 303 K with 600 mM KCl, and were individually replicated five times. *N_TOTAL_*: the number of all atoms, *N_POT_/Cl_CL_*: the number of K^+^ and Cl^-^, *N_WAT_*: the number of water molecules, *N_POPC_*: the number of POPC, *Dim*: X/Y/Z box dimension (nm).

| | $N_{TOTAL}$ | $N_{POT/CL}$ | $N_{WAT}$ | $N_{POPC}$ | $Dim$ |
| --- | --- | --- | --- | --- | --- |
| **Dark-state** | 261908 | 626/614 | 56824 | 528 | 10/10/25.6 |
| **Light-act.** | 262706 | 630/614 | 57054 | 530 | 10/10/25.6 |
| **Open state** | 262786 | 636/612 | 57082 | 530 | 10/10/25.6 |

**Supplementary Table 4:** Applied positional/dihedral angular restraints during energy minimization, equilibration simulations of C1C2. Time step: *Δt* (fs), individual simulation time: *t* (ns), force constant (kJ/mol/nm) for position/angle harmornic restraints of backbone heavy atoms (*k_bb_*), side-chain heavy atoms (*k_sc_*), lipid head group (*k_head_*), and lipid chirality and cis double bond (*k_torsion_*).

| | $\Delta t$ | $t$ | $k_{bb}$ | $k_{sc}$ | $k_{head}$ | $k_{torsion}$ |
| --- | --- | --- | --- | --- | --- | --- |
| Energy minimization | - | - | 4000 | 2000 | 1000 | 1000 |
| Equilibration 1 | 1 | 0.25 | 4000 | 2000 | 1000 | 1000 |
| Equilibration 2 | 1 | 0.25 | 2000 | 1000 | 400 | 400 |
| Equilibration 3 | 1 | 0.25 | 1000 | 500 | 400 | 200 |
| Equilibration 4 | 2 | 1 | 500 | 200 | 200 | 200 |
| Equilibration 5 | 2 | 1 | 200 | 50 | 40 | 100 |
| Equilibration 6 | 2 | 1 | 50 | 0 | 0 | 0 |
| Equilibration 7 | 2 | 100 | 0 | 0 | 0 | 0 |

**Supplementary Table 5:** Ion conduction derived from the MD-based CompEL simulations. . *Δq*: charge imbalance (*e*), *V*: membrane potential (mV), *N_I_*: total inward permeations, *G_I_*: inward conductance (pS), *N_O_*: total outward permeations, *G_O_*: outward conductance. All simulations were individually replicated five times at 303 K with ion concentration of 600 mM KCl.

| | $\Delta q$ | $t$ | $V$ | $N_I$ | $G_I$ | $N_O$ | $G_O$ |
| --- | --- | --- | --- | --- | --- | --- | --- |
| **Dark-state** | 4 | 1 | 343±36 | 0 | 0 | 0 | 0 |
|  | 6 | 2 | 503±30 | 0 | 0 | 0 | 0 |
| **Light-act.** | 6 | 2 | 531±26 | 0 | 0 | 0 | 0 |
| **Open state** | 6 | 2 | 389±27 | 28 | 1.2±1.1 | 4 | 0.2±0.4 |

## Notes

### Competing Interest Statement

The authors have declared no competing interest.

